# Soluble ECM promotes organotypic formation in lung alveolar model

**DOI:** 10.1101/2021.09.16.460165

**Authors:** Jonard C. Valdoz, Nicholas A. Franks, Collin G. Cribbs, Dallin J. Jacobs, Ethan L. Dodson, Connor J. Knight, P. Daniel Poulson, Seth R. Garfield, Benjamin C. Johnson, Brandon M. Hemeyer, Miranda T. Sudo, Jordan A. Saunooke, Braden C. Kartchner, Aubrianna Saxton, Mary L. Vallecillo-Zuniga, Matheus Santos, Brandon Chamberlain, Kenneth A. Christensen, Greg P. Nordin, A. Sampath Narayanan, Ganesh Raghu, Pam M. Van Ry

## Abstract

Micropatterned suspension culture creates consistently sized and shaped cell aggregates but has not produced organotypic structures from stable cells, thus restricting its use in accurate disease modeling. Here, we show that organotypic structure is achieved in hybrid suspension culture via supplementation of soluble extracellular matrix (ECM). We created a viable lung organoid from epithelial, endothelial, and fibroblast human stable cell lines in suspension culture. We demonstrate the importance of soluble ECM in organotypic patterning with the emergence of lumen-like structures with airspace showing feasible gas exchange units, formation of branching, perfusable vasculature, and long-term 70-day maintenance of lumen structure. Our results show a dependent relationship between enhanced fibronectin fibril assembly and the incorporation of ECM in the organoid. We successfully applied this technology in modeling lung fibrosis via bleomycin induction and test a potential antifibrotic drug *in vitro* while maintaining fundamental cell-cell interactions in lung tissue. Our human fluorescent lung organoid (hFLO) model represents features of pulmonary fibrosis which were ameliorated by fasudil treatment. We also demonstrate a 3D culture method with potential of creating organoids from mature cells, thus opening avenues for disease modeling and regenerative medicine, enhancing understanding of lung cell biology in health and lung disease.

## Introduction

The extracellular matrix (ECM) is classically known for its roles in cell anchorage and mechanical signaling in organs.[1] The role of ECM as an anchor is important in modulating organogenesis,[2, 3] cell differentiation,[4] cell survival,[5] and metabolism.[6] These ECM characteristics have been exploited by tissue engineers to recreate tissue- and organ-like 3-dimensional (3D) biological materials essential for disease modeling[7, 8] and regenerative medicine.[9] The tunability of ECM stiffness affects stem cell differentiation.[10] However, the roles of ECM as a soluble supplement are rarely studied and not well defined.[11] The majority of 3D culture techniques can be broken down into two categories: solid scaffold using some type of ECM or ECM-like synthetic materials and scaffold free, no ECM-like scaffolds.[12, 13] Both culture techniques have advantages and disadvantages.

In tissue engineering, the use of gelled ECM or other solid ECM-like polymers are prevalent in the construction of organotypic structures.[12] Scaffold-based cultures mimic the presence of a solid ECM akin to that of native organ tissue.[14] By this, the cells are provided with a substrate for organotypic attachment leading to biochemical signaling.[1], The ECM has both constructive and instructive roles in tissues and organs. In several stem cell-based approaches, cell clusters are embedded in solid ECM scaffolds to encourage organotypic formation.[15] Some obstacles in solid scaffold-based cultures include slow formation of tissues, inconsistent growth of aggregates, difficult recovery, and high contamination for downstream analyses.[9, 12]

Scaffold-free techniques, on the other hand, often use forced aggregation of cells on ultra-low attachment surfaces and rely on self-patterning of tissue to create consistently sized, reproducible multicellular aggregates.[9] Due to high initial local cell concentration, self-assembly into tissue-like structures is relatively faster than scaffold-based methods.[12] This characteristic gives scaffold-free technique an edge for being used in downstream high-throughput experiments. However, the use of scaffold-free force aggregated cultures is limited to creating tissue-like structures. The lack of organotypic structures is attributed to lack of organ-like ECM during self-patterning.[16]

Until now researchers have struggled to produce organotypic models from stable cell lines. There are several definitions given for organoids in the literature.[17] The prevailing definition focuses on stem cell differentiation and self-patterning of stem-derived cells while embedded in ECM with the idea that stable and mature cells lines do not self-pattern as well into organ-like structures nor give rise to other cell types. However, we have used A549 cells because earlier studies have proven that epithelial cells such as the A549 can form organotypic aggregate structures with a central lumen when embedded in ECM.[18-20] In this article, we define an organoid as an organotypic 3D cell aggregate composed of multiple specialized cell types, luminal self-patterning, and extracellular matrix which mimic organ structure and function.[21] Our hypothesis is that incorporating non-gelling concentrations of ECM proteins will improve the currently used scaffold-free, suspension culture resulting in a hybrid culture method. This research demonstrates the importance of the ECM in promoting organotypic patterning, growth, and survival as soluble supplements in suspension culture. Previous studies show that the composite properties of ECM are capable of determining stem cell fate.[22] Additionally, soluble ECM enhances cell proliferation and survival in 2D mesenchymal cell cultures, and enhanced formation of tight junctions in hepatic spheroid cultures.[11] However, the effects of low, soluble ECM concentrations in organotypic self-patterning and growth have yet to be fully elucidated. We illustrate the use of soluble ECM in the development of an organotypic model from stable cells within a 14-day time period. We refer to it as human fluorescent lung organoid (hFLO). hFLO has airspace-like lumen-gas exchange units and a perfusable vasculature, not present in traditional scaffold-free spheroids.

Here, we provide two potential applications of the hFLO: modeling bleomycin-induced pulmonary fibrosis with a pre-clinical therapeutic and a hypoxic angiogenesis study. Pulmonary fibrosis (PF) in humans is a progressive manifestation of interstitial lung diseases of known and unknown cause and leads to distorted lung parenchyma, irreversible fibrosis, honeycomb lung, respiratory failure and death.[23] Major advances in the pathogenesis of human PF have been made utilizing *in vitro* and *in vivo* models of experimentally induced lung injury and PF. However, there is an unmet need to understand the pathogenesis of PF in human lung cells and tissue via 3D modeling. The creation of such models will result in needed therapeutic development and validation in the targeting the cascades of inflammation and fibrosis. While several agents are being investigated in the clinic to determine safety and efficacy of pharmacologic agents (clinicaltrials.gov), only two drugs have been approved for clinical use.

These antifibrotic agents, nintedanib and pirfenidone are currently used in clinical practice but merely slow down disease process.[24] A long standing and well accepted model used to understand pathogenesis and modulation of PF is the bleomycin induced injury model.^29^ Here, we show that with bleomycin induction, the hFLO is a viable model in mimicking PF features *in vitro* and could be applied in a preclinical drug trial of an antifibrotic drug.

## Materials and methods

### Media formulations

The media formulations used in this paper were outlined in Table S4.

### 2D cell culture and cell line development

The cell lines used in this paper are HFL1 (ATCC CCL-153), EA.hy926 (ATCCRL-2922), A549 (ATCC CCL-185), and Lenti-X™ 293T (632180, Takara Bio, CA, USA). These cells were maintained in 2D culture using the complete 2D growth media. 0.25% trypsin-EDTA (90057, Thermo Fisher Scientific, MA, USA) was used during subculture and split ratios were kept being within the suggested range by the manufacturer. Subculture was done every 2-3 days. Fluorescent tags were introduced using transfer plasmids that were kindly gifted by Pantelis Tsoulfas via Addgene. Specifically, we used pLV-mCherry (36084, Addgene, MA, USA), pLV-Azurite (36086, Addgene, MA, USA), or pLV-eGFP (36083, Addgene, MA, USA) with second-generation lentiviral vectors pMD2.G (12259, Addgene, MA, USA) and psPax2 (12260, Addgene, MA, USA) which were gifted by Didier Trono through Addgene. Viral gene vectors were transfected using PEI 25K (23966, Polysciences, PA, USA) into Lenti-X™ 293T cells. Lentiviral particles for eGFP, Azurite, and mCherry gene insertions were collected with the media and were transduced into A549, EAhy or HFL1, respectively using polybrene (TR-1003-G, Sigma-Aldrich, MO, USA) for 72 hr. Cell purification was then done using fluorescence-activated cell sorting (FACS).

### Fluorescence-activated cell sorting (FACS)

Cells and cell aggregates were dissociated using Accumax (AM105.500, Innovative Cell Technologies, CA, USA) for 10 minutes and 30 minutes, respectively and strained using a 100 µm cell strainer (15-1100-1, Biologix Research Company, MO, USA) twice. The cells were then resuspended in 1% Bovine serum albumin (BSA, ab41899, Abcam, Cambridge, UK)-PBS buffer. All the FACS experiments were performed using BDAria Fusion Special Order System (BD Biosciences, CA, USA) equipped with 355nm, 488nm, 568nm, and 640nm lasers. Single cells were gated using forward scatter area (FSC-A) versus forward scatter width (FSC-W) and then with side scatter area (SSC-A) versus side scatter width (SSC-W). Positive expressing cells were gated from a negative control. Cells were sorted into 2D growth media using an 85µm nozzle and a 0.32.16 purity mask. In depleting NCadherin^high^ A549 cells, dissociated cells were stained for an hour on ice with NCadherin antibody (1:200) conjugated in-house using an Alexa Fluor 647 conjugation kit (A32733, Thermo Fisher Scientific, MA, USA). The bottom 30% and top 10% of cells based on AF647 signal were then sorted to deplete or enrich for N-cadherin markers, respectively. Sorted cells were grown in 2D culture as described above. Sorting efficiency during purification of fluorescent cells was verified using an Amnis imaging cytometer and confocal microscopy.

### Ultra-low attachment (ULA) plate

Agarose (BP160-100, Thermo Fisher Scientific, MA, USA) was dissolved in DMEM/F12 media to a concentration of 2% and slowly boiled using a microwave until all particles are dissolved. 50 µL of the agarose solution was then transferred over to each well of the 96 well U bottom plate (10062-900, VWR, PA, USA) then let solidify at room temperature (RT). These plates were then UV irradiated for at least 30 min and kept in a 4°C refrigerator for storage for a maximum of 1 month. For a high-output culture, we used our novel 217-well 3D agarose silicone cast as described previously.[25]

### 3D cell culture

To calculate the seeding ratios for each cell line, literature search led us to lung alveolar cell ratios to be around 24% epithelial, 30% endothelial, and 37% interstitial.[26] By this, we simplified the seeding ratio to be roughly 1:2:2 (A549 epithelial: EAhy endothelial: HFL1 fibroblast). For suspension-based cultures, a seeding ratio of 1.0×10^4^ to 1.0×10^5^ total cells per aggregate was used and was divided to each of the cell line using the ratio above. 100-200 µL of mixed cell suspension in 3D media or ECM supplemented 3D media were seeded into each well of the 96-well ULA plate. The triculture suspensions were permitted to self-aggregate and self-organize with the help of gravity and centrifugation for 5 minutes at a maximum of 250xg For the first 2-3 days of culture, 10 µM Y-27632 ROCK inhibitor (10005583, Cayman Chemical Company, MI, USA) was supplemented to prevent dissociation-dependent cell death.[27] The cultures were then kept in a 37°C humidified incubator at 5% CO_2_. Media was refreshed every 2-3 days and cultures were kept for 14 days unless otherwise mentioned in-text.

For the gel scaffolded cultures, 8.0×10^4^ cells mL^-1^ in the same ratio were suspended in pre-gelled solutions of 3 mg mL^-1^ collagen I, 3 mg mL^-1^ Matrigel or 4 mg mL^-1^ Matrigel. Rat tail collagen I (C3867-1VL, Sigma-Aldrich, MO, USA) was diluted in DMEM/F12 media with phenol red and 1N NaOH was added dropwise until pH was around 7.4. Matrigel (354263, Corning, NY, USA) was simply diluted in DMEM/F12. The solutions were kept on ice to prevent gelling. Cells were resuspended in the pre-gelled solutions and 300 µL of the cell solutions were dropped into glass-bottom plates (P35G-1.5-14-C, MatTek, MA, USA). These were then gelled for 30 minutes in a 37°C humidified incubator, after which complete growth media was added to each plate and incubated. Media was refreshed every 2-3 days of culture.

### 3D aggregate growth assays

Triculture cell aggregates were grown for 14 days using either traditional force aggregation (FA) or as supplemented with 300 µg mL^-1^ Matrigel. ROCK inhibitor Y27632 was supplemented for the first 3 days of culture. The media was then refreshed every 2-3 days and the cell aggregates were photographed using either Amscope phase-contrast microscope (Amscope, CA, USA) or Thermo Evos XL phase-contrast microscope (Thermo Fisher Scientific, MA, USA). To measure their respective areas, images were imported into ImageJ as 8-bit format, after which a threshold was manually applied. The image was then processed five times iteratively with the dilate function to converge cells. Cell aggregate sections were then selected using the freeform selection tool and measured for area. The measurements were then normalized to the day 1 measurements. Line graphs were created using the seaborn library in Python.

### Multispectral imaging flow cytometry (MIFC)

Cells were dissociated using Accumax (AM105.500, Innovative Cell Technologies, CA, USA) for 10 minutes for 2D culture and 20 minutes for 3D cultures. These were then fixed using 1% paraformaldehyde (PFA) in PBS for 30 minutes at 4°C. Cells were then permeabilized using 0.1%Triton X-100 for 10 minutes and blocked using Seablock blocking buffer (37527, Thermo Fisher Scientific, MA, USA) for 30 minutes at 4°C. Primary antibodies were then diluted with 10% Seablock in 0.1% Tween20 and PBS (PBST) and incubated overnight at 4°C. Secondary antibodies were then added for 1h at 4°C. The samples were run in 1% BSA-PBS solution. Positive controls were run first to optimize laser power for each experiment. Using the raw max pixel data for each channel, laser power was adjusted to prevent signal saturation. Data was collected after gating out cells using phase contrast area vs aspect ratio. Both FCS and RIF files were collected for most of the experiments. Color compensation was done during analysis in IDEAS software (Amnis, WA, USA, version 6.2) using the compensation wizard tool, after which in-focus cells were gated using gradient RMS data. For the cell death-related morphometric features, we used bright-field circularity, aspect ratio, shape ratio, contrast, and modulation. Then, the expression values were exported into either FCS or TXT files. FCS files were plotted using python. Representative images were exported as TIFF and colorized either in IDEAS or in Adobe Photoshop 2020 (Adobe, CA, USA, version 21.0.2).

### Localization of Matrigel

A 10 mg mL^-1^ solution of Matrigel was conjugated with 20x molar excess concentration NHS-dPEG-biotin (QBD10200-50MG, Sigma-Aldrich, MO, USA) by mixing overnight at 4°C. Excess biotin was removed with a 5K MWCO concentrator (CLS431487-12EA, Millipore Sigma, MA, USA) at 4000xg for 100 minutes at 4°C. Matrigel concentration was then assayed with a BCA array (23225, Thermo Fisher Scientific, MA, USA) and diluted to 300 µg mL^-1^ in 3D media. To confirm biotinylation, 50µL of 3D growth media, hFLO media, and biotinylated hFLO media were loaded into 16% SDS-PAGE gel and transferred onto 0.2 μm nitrocellulose membranes (1620150, Bio-Rad, CA, USA) through electro blotting. The blot was then probed using Streptavidin Dy-Light 800 (21851, Invitrogen, 1:10,000, MA, USA) for an hour at RT. The blots were imaged using the Odyssey CLx (9140, Li-Cor, NE, USA).

To determine if the Matrigel localizes in the aggregate, 1.0 × 10^4^ cells per aggregate were cultured in a multi-well agarose dish as described above. For the first 24 h, normal hFLO media was used and this was changed into the biotinylated hFLO media after 24 h. Control aggregates were kept in hFLO media. These were then kept in culture for 48 more hours. After 3D culture, the cell aggregates were processed and imaged as described in the whole-mount immunofluorescence methods using Streptavidin (29072, Biotium, 1:500, CA, USA) and a DAPI (1:500) counter-stain.

### Flow Cytometry

Cells were dissociated using Accumax for 10 minutes for 2D culture and 20 minutes for 3D cultures. These were then fixed using 4% PFA in PBS for 15 minutes at 25°C. Cells were then permeabilized-blocked using 0.1%Triton X-100 in Seablock buffer for 15 minutes at 25°C. Primary antibodies were then diluted with 10% Seablock in 0.1% Tween20 and PBS (PBST) and incubated for 1h at 25°C. Secondary antibodies (1:500 dilution) were then added for 30 min at 25°C. The samples were run in 1% BSA-PBS solution. Detector voltages were determined using single-color controls. All files were exported as FCS and analyzed using FlowJo (BD, CA, USA, v9.9.4). Compensation was done using the Compensation Wizard and single-color controls. Doublets were then excluded using FSC-A vs FSC-H and SSC-A vs SSC-W gates. Single cells were then analyzed and any applicable gates are included in supplementary information.

### Murine and porcine lungs

All experiments were performed per the NIH Guide for the Care and Use of Laboratory Animals and were approved by Institutional Animal Care and Use Committees at Brigham Young University-Provo. Wild type C57BL/6 mice were sacrificed and were perfused ventricularly with sodium heparin solution in PBS (H3393-100KU, Sigma-Aldrich, MO, USA). The lungs were then excised and fixed in 4% PFA overnight at 4°C. The porcine lungs were gifted by Circle V Meats Co. (UT, USA). The lungs were received and processed within the hour of sacrifice. Lung sections were washed thrice with heparinized PBS for 10 min each and visible blood clots were removed. The lungs were then sectioned into 1-inch cubes and fixed overnight with 4% PFA at 4°C.

### Live confocal imaging

Suspension-based cell aggregates were collected and transferred to a MatTek glass bottom plate (P35G-1.5-14-C, MatTek, MA, USA) with minimal volume of complete cell media to prevent dehydration. The plates were then mounted into the live cell imaging chamber (Okolab, Ottaviano, Italy) attached to a Leica TCS SP8 confocal microscope (Leica, Wetzlar, Germany). The chamber was set to 5% CO_2_ at 37°C throughout the experiment. Either the 20X or 40X objectives were used during imaging. Argon laser was consistently set to 25% and individual laser powers were kept being less than 5% to prevent excessive phototoxicity while imaging. Images were acquired with at least 512×512 voxel density with optical zoom from 0.7 to 1.0X.

### Section and 3D whole-mount immunofluorescence

Samples were fixed in 1% PFA overnight at 4°C 800rpm shaking incubator, embedded in Tissue Tek OCT (4583, Sakura, Tokyo, Japan), and then sectioned into 7-10 µm sections using a TN50 cryostat (TNR TN50, Tanner Scientific, FL, USA). The slides were dried for 15 minutes at room temperature prior to storage in -80°C or processing for immunohistochemistry or histology. The sections were then permeabilized using 0.1% Triton X-100 for 10 minutes and then blocked using Seablock buffer for 1hr at room temperature. Primary antibodies (Table S4) were diluted in 10% Seablock in PBST and incubated overnight at 4°C. After intensive washing, secondary antibodies were then diluted to 1:500 in 10% Seablock PBST and incubated for an hour at RT. Counterstaining for the nucleus, if necessary, was done either by DAPI (14285, Cayman, MI, USA; 1:500), Hoecsht (62249, Thermo Fisher Scientific, MA, USA; 1:1000), or Toto-3 iodide (T3604, Invitrogen, MA, USA; 1:500) diluted in PBST. The slides were mounted using diamond antifade (P36961, Thermo Fisher Scientific, MA, USA) and cure at RT for at least 24h prior to imaging.

For whole mount imaging, each step was done in an 800-rpm temperature-controlled shaker. Permeabilization was achieved using 0.3% TritonX-100 for 15 minutes then blocked for 2h at RT using Seablock blocking buffer with 0.3%TritonX-100. Primary antibodies were diluted in 10% Seablock in 0.3% Triton X-100 and incubated for 24-48h at 4°C. Secondary antibodies and nuclear counterstain were added to the whole aggregates for 2h at RT. After the samples were immunostained, refractive index matching was achieved using an in-house Ce3D clearing buffer overnight at RT as previously shown.[28]

Confocal imaging was done using a Leica TCS SP8 Hyvolution confocal microscope with LASX software (Leica, Wetzlar, Germany, version 3.1.1.15751). Argon laser power was consistently set to 25%. Use of the HyD detectors were preferred over the PMT detectors and gains for the detectors are set to less than 100%. If needed, samples were stitched using the built-in Auto Stitch function in the LASX software. The images were deconvolved Huygens Essential (SVI, Hilversum, The Netherlands, version 19.04) using the Deconvolution Wizard function. The final images were exported as TIFF files and gamma was not adjusted for any images. Volumetric image analysis was perfomed using the 3D analysis package in LASX software. Volumes were then normalized to nuclear volume.

### Histological staining

#### Hematoxylin and Eosin (H&E) staining

H&E staining was performed on cell aggregate frozen sections as indicated by the manufacturer (Richard-Allen Scientific, MI, USA), using hematoxylin (7211, Richard-Allen Scientific, MI, USA) and eosin staining (7111, Richard-Allen Scientific, MI, USA). The slides were then cleared 3 times in 100% ethanol. Slides were then washed 3 times using Histo-Clear (HS-200, National Diagnostics, NC, USA) for 5 minutes, after which the slides were mounted using Clarion mounting medium (CO487, Sigma-Aldrich, MO, USA). Slides were then cured overnight at 37°C before imaging.

#### Masson staining

Cell aggregate sections were processed following the protocol of Trichrome Stain (Masson) as indicated by the manufacturer (HT15-1KT, Sigma-Aldrich, MO, USA), after which the slides were allowed to air dry. Slides were then washed using Histo-Clear (HS-200, National Diagnostics, NC, USA) for 5 minutes before being mounted using Clarion mounting medium (CO487, Sigma-Aldrich, MO, USA). Slides were then cured overnight at 37°C before imaging.

#### Hematoxylin-DAB

Cell aggregate sections were permeabilized using 0.1% Triton X-100 for 10 minutes and then blocked using Seablock buffer for 1hr at room temperature. Primary antibodies (Table S4) were diluted in 10% Seablock in PBST and incubated overnight at 4°C. After intensive washing, HRP-conjugated secondary antibodies were then diluted to 1:500 in 10% Seablock PBST and incubated for an hour at RT. The chromogen was developed using a DAB kit (34002, Thermo Scientific, MO, USA) for 15 min at RT. Counterstaining for the nucleus was done using hematoxylin staining and bluing reagent. The slides were then cleared 3 times in 100% ethanol. Slides were then washed 3 times using Histo-Clear (HS-200, National Diagnostics, NC, USA) for 5 minutes, after which the slides were mounted using Clarion mounting medium (CO487, Sigma Aldrich, MO, USA). Slides were then cured overnight at 37°C before imaging.

### Transmission electron microscopy (TEM)

Samples were collected directly into 2% Glutaraldehyde in 0.06 M sodium cacodylate buffer and remained in the fixative for at least 2 hours. The samples were washed 6 times in PBS for 10 minutes each. Secondary fixation continued using 1% Osmium tetroxide (19152, Electron Microscopy Sciences, PA, USA) for 1.5 hours. Samples were then washed 6 times with distilled water for 10 minutes each. The samples were then stained with 0.5% Uranyl Acetate overnight with gyration. In preparation for resin embedding the samples were dehydrated serially with ethanol (30%-50%-75%-90%-100%-100%-100%) with 10-minute incubation per stage. Samples then were washed 3 times in acetone for 10 minutes each. Resin embedding was done using in house Spurr low viscosity resin with serial preparation steps in acetone (2:1-1:2-0:1, acetone:resin) for 90 min each with gyration.[29] The samples were then resin-embedded in Beem capsules (130, Ted Pella, CA, USA) and cured overnight in a 70°C oven. The embedded samples were then sectioned to 80 nm thick using an RMC MTX microtome using an in-house glass knife. Sections were collected on a Formvar/Carbon 200 mesh support grids (01800, Ted Pella, CA, USA) and counterstained with lead. Images were taken using a Helios NanoLab 600 Dual Beam FIB/SEM using the TEM feature.

### Shotgun proteomics

Fourteen-day old FA and hFLO cell aggregates were collected and washed twice with PBS for 10 minutes each with gyration. To lyse the samples, RIPA buffer with Halt’s inhibitor cocktail (89900, Thermo Fisher Scientific, MA, USA) was added and then the samples were vortexed thrice for 5 minutes each time and then boiled for 5 minutes at 95°C. The samples were then further sheared using a bead beater and then passed through a 25G needle. The whole aggregate lysates were collected by centrifugation at 16,000xg for 5 minutes. Protein concentration was then quantified using Peirce™ BCA Protein assay kit (23225, Thermo Fisher Scientific, MA, USA). Processing the samples for proteomics was then continued using a 10 kDa filter aided sample preparation protocol previously published with minor modifications.[30] The proteins were digested using Lys-C enzyme (P8109S, New England Biolabs, MA, USA; 1:50) and then with trypsin (V5111, Promega, WI, USA; 1:100). The samples were then dried in a vacuum centrifuge for 1.5h at RT and then resuspended in running buffer. The samples were then analyzed using Thermo Fisher™ Q-Exactive Obitrap (Thermo Fisher Scientific, MA, USA). The mass spectrometry proteomics data have been deposited to the ProteomeXchange Consortium via the PRIDE[31] partner repository with the dataset identifier PXD027296.

### Proteomic bioinformatic analysis

Mass spectrometric raw data files were processed using PEAKS (Bioinformatics Solutions, Waterloo, Canada, version X PRO 2020) and were searched against the human proteome database (Uniprot_SwissProt_Oct2020). The Label-free quantification (LFQ) values were quantile normalized between FA and hFLO proteome. LFQ values were compared between the FA and hFLO proteomes using pairwise statistical tests based on two-tailed Welch’s t-test. Protein-protein correlation was calculated using R Studio corrplot function.[32] Unbiased gene set interrogation was performed using GSEA native GO-term set (ftp://broadinstitute.org://pub/gsea/gene_sets/c5.all.v7.4.symbols.gmt).[33] Graphs were then created using GraphPad Prism (GraphPad Software, CA, USA) or R Studio ggplot2.

### Bleomycin-induced fibrosis

Cell aggregates were grown as described above for 7 days, after which the cell aggregates were treated with 20 µg mL^-1^ bleomycin (13877, Cayman Chemical Company, MI, USA, stock solution diluted in either PBS or DMSO). Media was changed after 3 days of culture with bleomycin (i.e., at day 10). At day 10, the organoids were weaned off the bleomycin insult and then treated with either 10µM fasudil (B3523, ApexBio, TX, USA) or with DMSO (control, vehicle used for fasudil). The cultures were kept in treatment media for 4 more days with media changing every two days. Cell aggregates and media were collected and fixed as described above.

### Cytokine Array

The organoid media from the three treatment groups were collected at day 14. The levels of cytokines secreted to the media were immediately evaluated using the Human Cytokine Array C1000 (AAH-CYT-1000-8, Ray Biotech, GA, USA) with minor modifications. Briefly, the membranes were blocked at room temperature for 30 minutes using the blocking buffer. The media from the organoids were transferred onto the membranes and incubated with gentle rocking overnight at 4°C. After washing, the membranes were incubated with the biotinylated antibody cocktail overnight at 4°C. After further washes, the membranes were probed with Streptavidin-CF680R (29072, Biotium, CA, USA 1:1000) for 1h at RT. The membranes were imaged using an Odyssey CLx imager (LI-COR, Cambridge, UK) at high resolution and a wavelength of 700 nm. The exported images of the membranes were quantified using gel analysis plugin on FIJI distribution of ImageJ (open source) under the GNU General Public License.

### Hypoxic culture

At day 7, media was changed to angiogenic media comprised as described in Table S4 and the culture was purged with 5% O_2_, 5% CO_2_, 90% N_2_ gas mixture. Cell media was changed every 2-3 days using the angiogenic media. The air inside the chamber was replaced every time media was changed. The culture was harvested 7 days post induction.

### 3D printing

The 3D printed high-flow organoid devices used in this paper were designed in OpenSCAD and fabricated using a custom stereolithographic 3D printer. The printer was a third-generation model of the printer that was previously published.[34] The printer’s custom Python software allows for precise control over all aspects of the printing process enabling intricate design capabilities, such as the grates used on the organoid devices. A custom photo-polymerizable resin was used consisting of poly(ethylene glycol) diacrylate (PEGDA, MW258, 475629, Sigma-Aldrich, MO, USA) as the monomer, 1% (w w^-1^) phenylbis(2,4,6-trimethylbenzoyl) phosphine oxide (Irgacure 819, 415952, Sigma-Aldrich, MO, USA) as the photo-initiator, and 0.38% (w w^- 1^) avobenzone (16633, Sigma-Aldrich, MO, USA)—a UV absorber to control light penetration during 3D printing. This was an ideal surface for organoid analysis because it has been proven to be non-cytotoxic with extremely low adherence.[35] The resulting device has the dimensions shown in Table S5.

### Millifluidic dextran perfusion

Cell aggregates including A549 spheroids, triculture aggregates grown in traditional FA, hFLO cell aggregates, and vhFLO cell aggregates were grown until 14 days post seeding. Capillary fluidic flow was modeled using PTFE tubing (AO-06417-21, Cole-Palmer, IL, USA), a peristaltic pump (61161-354, VWR, PA, USA), and a 3D-printed biocompatible 4-well chip based on our previously published methodology.[35] Prior to the experiment, the fluidic system was flushed with DMEM/F12 media and the peristaltic pump was then calibrated manually to 1 mL min^-1^well^-1^. Afterwards, 20 μg mL^-1^ Texas Red™ 70 kDa dextran (D1864, Invitrogen, MA, USA) was perfused through the system to replace the media. 14-day old cell aggregates were then transferred into the 3D-printed microwell devices. To seal high-flow devices, glass slides wrapped with parafilm were carefully clamped onto the 3D printed chips. Dextran solution was perfused cyclically for one hour, after which cell aggregates were collected and rinsed with PBS thrice for 5 minute each to remove unbound dextran. They were then fixed in 1% PFA overnight at 4°C. The cell aggregates were then counterstained with DAPI and processed for whole-mount immunofluorescence as described above.

### Image analyses

#### Line profile grey value analysis

Day 14 cell aggregate images were uploaded to ImageJ version 1.53c. We used the line tool and the plot profile function to derive grey values (Y) versus the line distance (X). We then used Microsoft Excel for further data processing. We normalized the grey values (Y) to a common scale where denser regions of tissue have the higher values (100%) and spaces/lumina have lower values (0%). The distance value (X) is normalized such that the core of the cell aggregate is set at zero (range= -100, +100). For more information, see supplementary protocol.

#### Lumen size quantification

Histologically stained samples were imaged and processed with ImageJ software. First, images were converted to an 8-bit greyscale, after which a threshold was applied. The image was then processed five times iteratively with the dilate function to converging boundary cells.

Afterwards, the section area was selected, the image outside the aggregate section was cleared, and the image was inverted such that the lumens only were displayed. The “count particles” function was used to measure lumen characteristics. Data was processed in Microsoft Excel, lumen under the size of 200 μm^2^ were excluded in all analysis except the calculations of total lumen area and lumen percent area.

#### Masson-Trichrome and H-DAB color analyses

The ratios of collagen to tissue in organoids were determined by analyzing images of Masson-Trichrome stained sections. These images were separated into red, blue, and green components using ImageJ using color deconvolution function as previously described.[36] For the Masson staining, areas of the blue and red image channels were determined by thresholding and collagen deposition was determined as the ratio of blue area to red area (collagen to tissue). Intensity histograms were created using the ImageJ histogram function on the selected red area, and average pixel intensity was calculated from the histogram output in Microsoft Excel. For H-DAB, the target protein is the brown channel and the hematoxylin is the blue-purple channel.

### Statistical analyses

Independent samples were randomly selected for all studies. Statistics were determined using GraphPad Prism (GraphPad Software, CA, USA, version 9.1.1). Where appropriate, equal sampling was employed. Pairwise comparisons to determine statistical significance was carried out using two-tailed Welch’s t-test. For multiple group comparisons, One-way ANOVA was employed with pairwise tests carried out with Welch’s correction. Graphs were then generated using GraphPad Prism 9, Python, or R Studio 3.6.2.

## Data Availability

The mass spectrometry proteomics data are publicly available. They have been deposited to ProteomeXchange Consortium via the PRIDE[31] partner repository and can be accessed with the dataset identifier PXD027296. All supporting the main and supplementary conclusions of the paper were provided with the manuscript. All other data are available upon reasonable request from the corresponding author, P.M.V.R.

## Results

### Soluble ECM promotes airspace-like lumen-gas exchange unit formation in suspension culture

Previous studies have used soluble matrix proteins as media additives to improve cell aggregation[37, 38] and as a potent pro-proliferative factor.[11] Despite this, the relationship between soluble ECM and self-patterning towards 3D organotypic growth is yet to be discovered. To model the lung alveolus, we used traditional forced aggregation (FA) method and our hybrid method supplementing soluble concentration of ECM to promote organotypic self-patterning. In order to visualize topographical changes accompanying ECM changes, three fluorescently labeled stable cell lines were employed to represent the three major alveolar tissue types including endothelial EA.hy926 (EAhy, Azurite-tagged), lung epithelial type II-like A549 (eGFP-tagged), and normal lung fibroblast HFL1 (mCherry-tagged) (Supplementary Fig. 1a-c). Previous studies show that A549 cells form organotypic luminal structures once embedded in ECM, mimicking features of an organotypic culture observed using primary-derived alveolar type 2 cells.[19, 39, 40] Moreover, A549 has previously been shown to exhibit stem-like properties of self-renewal, multipotency, differentiation, and clonogenic formation in its epithelial (EPCAM+) subpopulation.[41-44] On the other hand, due to its nature as a cancer cell line, certain subpopulations express mesenchymal, pro-metastatic proteins.[45] To mitigate these traits and improve its epithelial stem-like nature, we used fluorescence-activated cell sorting (FACS) to deplete mesenchymal N-Cadherin^high^ population which selects for the more stable epithelial cell population.[46]

**Fig. 1:**
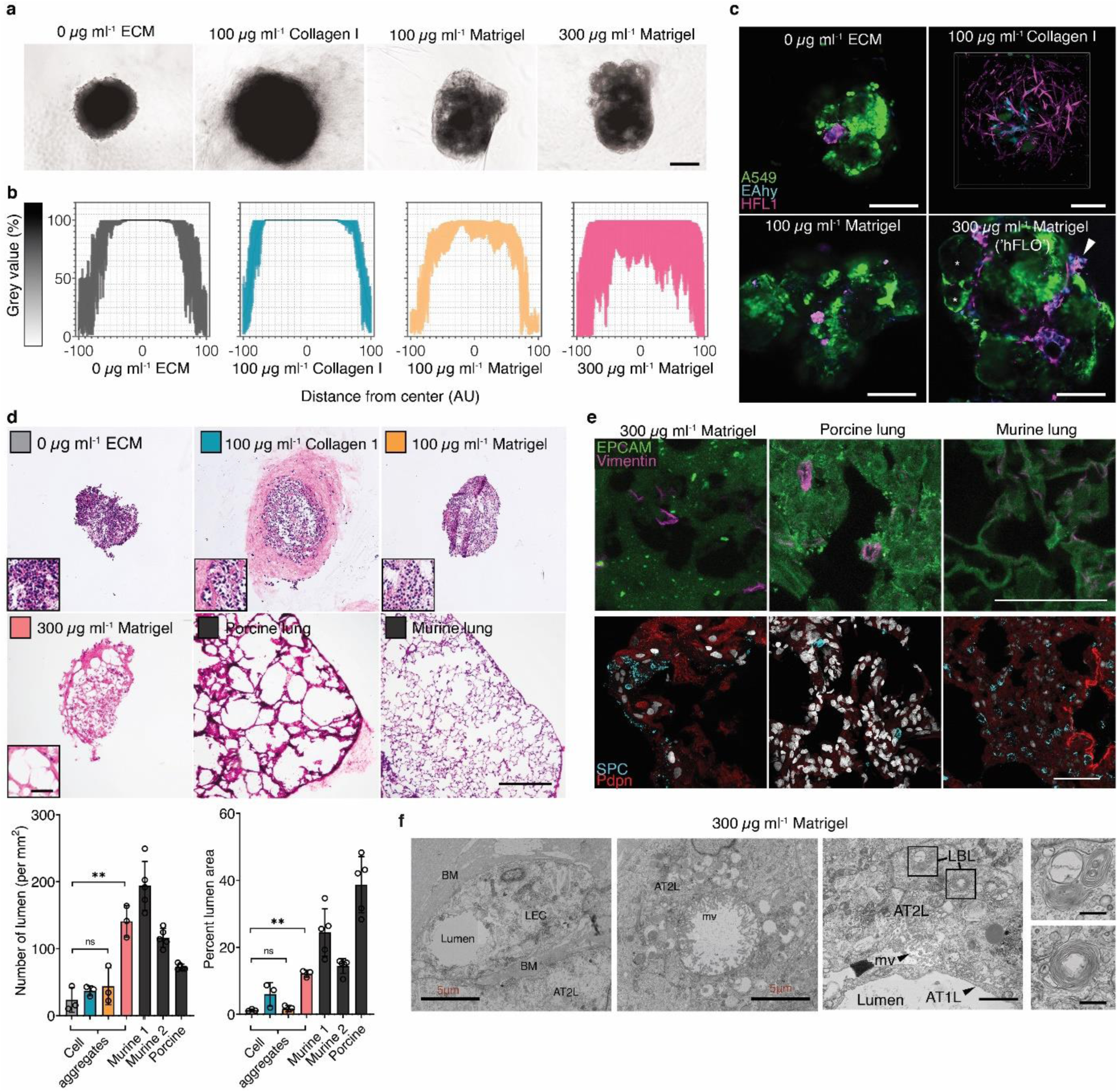
Alveolar-like lumen structure formation. **a**, Representative bright-field images of 3D cell aggregates after 14 days of culture. Scale bars, 300 µm. **b**, Effect of soluble ECM in aggregate density. Line profile analysis at the major axis of the aggregates expressed as normalized grey value. Lines indicate the grey value range at the specific normalized distance from the center. **c**, Live 3D fluorescence confocal image (single z-section) showing the self-organized localization of the three cell lines—A549 (green), EAHy (blue), and HFL1 (magenta). In aggregates grown using 300 µg/mL Matrigel, organotypic structures such as the lumina (*) and endothelial networks (arrow) are indicated. Scale bars, 200 µm. **d**, Hematoxylin and Eosin staining of the aggregates showing that the lumen formation in 300 µg/mL Matrigel aggregates is comparable to the mammalian lung parenchyma. Scale bars, 300 µm for full image; 50 µm for inset. Lumen formation is quantified using number of lumen normalized to aggregate size and percent lumen area. Bar colors are indicated in the micrographs above. Mean, individual measurements, and standard deviation are shown. Three independent cell aggregates and lungs from two mice and one pig were used. \*\**P*<0.01, ns (*P*>0.05) determined by One-way ANOVA with Welch’s correction. **e**, Fluorescence confocal images (maximum projection) showing localization of epithelial cells (EPCAM, green) and mesenchymal cells (Vimentin, magenta) relative to the lumen. Specific alveolar epithelial subtypes: Type I (Podoplanin, red) and Type II (prosurfactant protein C, blue). Scale bars, 50 µm. **f**, Transmission electron microscopy of cell aggregates showing stratification and lumen formation in 300 µg/mL Matrigel aggregates. Alveolar type II-like epithelial cells (AT2L) shown with their lamellar body-like inclusions (LBL) and microvilli (mv). Other features include: thin, alveolar type I-like cells (AT1L), a luminal structure, presence of a basement membrane (BM), and luminal endothelial cells (LEC). Scale bars, 3 µm for full images; 500 nm for zoomed images.

We then engineered a 3D triculture to model the lung alveolus and the morphological changes accompanying soluble ECM concentration. In the native lung alveolus, the epithelium forms airspaces with interstitial and vascular cells between the epithelium. We incorporated labeled or unlabeled A549-eGFP, HFL1-mCherry, and EAhy-Azurite in 3D tricultures with varying ECM concentrations—high (gel scaffold, ≥3 mg ml^-1^), low (soluble ECM-supplemented suspension, ≤300 µg ml^-1^), and no ECM (traditional FA) (**Fig. 1** and Supplementary Fig. 2). The 3 cells were mixed in suspension prior to seeding and spheroid organization was dependent on self-patterning aided by soluble ECM. Cell seeding ratios were followed based on endogenous human lung cell composition which we determined to be roughly 1:2:2 (epithelial: interstitial: vascular) (Table S1).[26] With this ratio, high ECM concentrations in gel scaffolds led to the formation of multiple A549 epithelial clusters without cohesion of all three cell lines (Supplementary Fig. 2a-b). In gel scaffolds, fibroblast-endothelial (HFL1-EAhy) network formation were observed as reported by previous researchers[47, 48] (Supplementary Fig. 2b). In comparison, all soluble ECM concentrations tested led to the formation of a singular cell cluster with all three cell lines incorporated (**Fig. 1a**). An important component of the human alveolus is alveolar sacs, which are small airspace-filled voids made from epithelial cell and separated by a common septum.

Bright-field images of cultures with soluble ECM were analyzed for possible less dense airspace-like voids. Images and grey line profile analysis show several less dense regions with supplementation of 300 µg ml^-1^ Matrigel, while traditional FA cultures appear as densely packed aggregates (**Fig. 1b**). Live confocal imaging reveals self-organized structures in the triculture aggregates similar to those observed in mammalian alveolus (**Fig. 1c** and Supplementary Fig. 2b-c). Self-organized structures observed in both gel scaffold and soluble ECM-supplemented cultures include epithelial clustering and an endothelial-fibroblast network, neither of which were observed in the traditional FA culture (0 µg mL^-1^) (**Fig. 1c** and Supplementary Fig. 2b-c, 3a-b). In soluble 100 µg ml^-1^ collagen suspension cultures, epithelial cells form a dense core aggregate; however, 300 µg ml^-1^ Matrigel cultures form airspace-like lumina (**Fig. 1a-c** and Supplementary Fig. 2c, 3a-b).

Aggregates were stained with hematoxylin and eosin (H&E) and compared to healthy murine and porcine lung sections to further confirm the relationship between the use of soluble ECM and the formation of voids inside the aggregates. Triculture aggregates grown with traditional FA (0 µg mL^-1^ ECM), 100µg mL^-1^ collagen I, and 100µg mL^-1^ Matrigel were confirmed to be predominantly compact cell clusters. Yet with 300 µg mL^-1^ Matrigel supplementation, we observed the presence of mammalian airspace-like lumina with septa (**Fig. 1d**). Upon quantification of the soluble ECM supplemented aggregates, we saw an increased number and percent area of voids inside the 300 µg mL^-1^ Matrigel-grown aggregates relative to the traditional FA. Together this data confirms that supplementation of soluble Matrigel leads to topographical arrangement of cells similar to those found in native alveoli (Supplementary Fig. 3).

Keeping our overarching goal in mind to produce an organotypic structure for use in downstream processes, we used established mammalian alveolus markers that are cell type-specific and are known to localize to specific regions in the alveolus. Immunohistochemistry of day 14 cell aggregates were compared to murine and porcine lung sections to verify the typical mammalian lung alveolus localization pattern. Epithelial cells as indicated by epithelial cell adhesion molecule (EPCAM+) are known to localize on the airspace-like lumen with mesenchymal cells labeled with (Vimentin+) in the alveolar-like septum (**Fig. 1e**). Epithelial subpopulation type II (pro-surfactant protein C, SPC+) and type I (Podoplanin, Pdpn+) cells should be found proximal to the lumina. Observational comparisons show that all four of these markers distributed in aggregates grown using 300 µg mL^-1^ Matrigel in a similar fashion to both murine and porcine sections. (**Fig. 1e**).

Transmission electron microscopy (TEM) was used to visualize vital ultrastructures in the aggregates such as the presence of lamellar bodies and microvilli in type II cells, presence of basement membrane, and the localization of the epithelial cells bordering the airspaces. We observed the presence of lamellar body-like (LBL) inclusions A549 (type II-derived cell line) as previously shown. We also identified LBL inclusions in 0 µg mL^-1^ ECM and 300 µg mL^-1^ Matrigel cell aggregates and A549 spheroids which is attributed to A549 being a type II-derived cell (**Fig. 1f** and Supplementary Fig. 4a-b). Using TEM, we verified that the use of 300 µg mL^-1^ Matrigel led to the formation of luminal structures—both on the alveolar epithelial surface and within the luminal endothelial cells (**Fig. 1f**). Moreover, we observed the presence of a basement membrane between epithelial and endothelial tissue layers in 300 µg mL^-1^ Matrigel cell aggregates further supporting observations of organotypic formation (**Fig. 1f**). Together, our data suggests that basic alveoli-like and vascular structure is achieved in a triculture suspension by supplementing with 300 µg mL^-1^ Matrigel. Hereafter, we will refer to this triculture hybrid 3D method as the human fluorescent lung organoid (hFLO) (Supplementary Figure 3a-b).

### Soluble ECM promotes aggregate growth and survival compared to FA cultures

Previous studies show pro-survival effects of a soluble ECM in mesenchymal stem cells.[11] Thus, we examined pro-survival and pro-growth effects of soluble Matrigel in 3D suspension. We tracked aggregate size using bright-field microscopy during a 14-day culture of both FA and hFLO and found that hFLO aggregates increase 2-fold in size while FA aggregates remained nearly constant (**Fig. 2a-b**). Dissociated aggregate cell counts revealed a 3-fold increase in cell counts in hFLO, whereas FA aggregates only increased 1.5-fold from original seeding density (**Fig. 2c**). To determine if supplementation of soluble ECM supports the maintenance of the general aggregate structure, we cultured hFLO up to day 70 and examined general histological structure using H&E staining. Although we saw a slight increase in percent lumen area, no change was observed in average number of luminal voids, suggesting that the general organotypic structure can be maintained for at least 70 days (**Fig. 2d**).

**Fig. 2:**
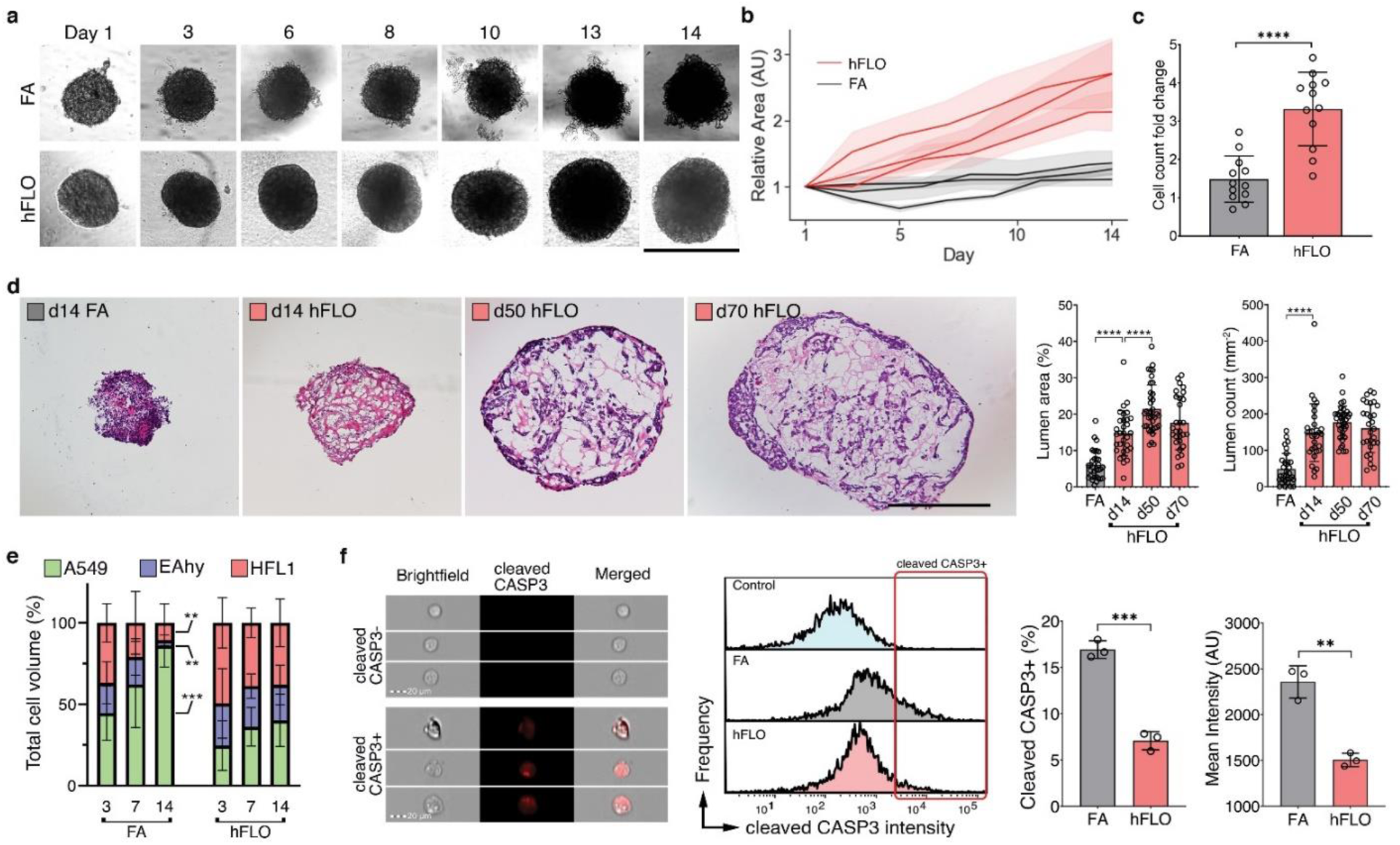
3D aggregate growth and cell survival. **a**, Bright-field images of 3D cell aggregates during the 14-day culture. Scale bar, 300 µm. **b**, Line graph showing change in aggregate area. Areas were normalized to their respective day 1 area. Means and standard deviation are shown. Individual line indicates independent experiments performed by independent researchers. N = 10 independently sampled aggregates on each measured day. **c**, Cell counts at day 14-fold change from day 0 seeding density. Means, individual values, and standard deviation are shown. 3 independent experiments were performed with >3 cell aggregates used; N=12 aggregates. \*\*\*\**P*<0.0001 determined by two-tailed t-test with Welch’s correction. **d**, Hematoxylin and Eosin staining of cell aggregates showing stability of lumen structure in hFLO (up to 70 days of culture) as quantified using lumen area and count. Mean, individual values, and standard deviation are shown. \*\*\*\**P*<0.0001 determined using two-tailed t-test with Welch’s correction using values from n=27 independent aggregates. **e**, Cell population composition at days 3, 7, and 14 of culture; n=7 independently sampled aggregates. \*\**P*<0.01, \*\*\**P*<0.001 determined using One-way ANOVA with Welch’s correction comparing day 14 to day 3 values. **f**, Expression of cleaved caspase 3 of day 14 cell aggregates using imaging flow cytometry. Three independent plates (around 200 cell aggregates) were used and an unstained control from pooled samples was used as a negative gating control. Left bar plot shows percent positive cells with mean, individual points, and standard deviation indicated. Right bar plot shows mean intensities found in each of the samples. \*\*\**P*<0.001 determined using two-tailed t-test with Welch’s correction.

One of the drawbacks of using FA culture is that within the dense aggregate core cell death occurs due to hypoxia and starvation.[49] Previous research demonstrates that endothelial cells are susceptible to cell death when cultured in suspension.[50] Moreover, in precision cut lung slice cultures, endothelial cells were observed to not survive beyond 7 days of culture.[51] Cell type survival was measured through volumetric analysis of each fluorescently-labeled population to determine cell ratio changes throughout the 14-day culture in suspension. Without soluble ECM, we found that HFL1-fibroblast and EAHy-endothelial cell populations progressively decrease through the time of culture with A549-epithelial cells constituting greater than 80% of FA by day 14. (**Fig. 2e**). In contrast, cell population ratios based on volume were maintained in hFLO for the same time period (**Fig. 2e**).

To determine if cell death was activated in FA more than in hFLO, we performed imaging flow cytometry. We found that apoptotic marker, cleaved caspase 3 (CASP3) is significantly increased in FA (**Fig. 2f**). Moreover, cell death morphometric analyses for shape change (cell circularity, aspect ratio, and shape ratio) show increased apoptotic markers in FA (low circularity, low aspect ratio, and low shape ratio, Supplementary Fig. 5).[52, 53] Our data suggests that hFLO with 300 µg mL^-1^ Matrigel supplementation resulted in enhanced aggregate growth with conserved cell populations and histological structure.

Our results imply that the supplementation of 300 µg mL^-1^ Matrigel is responsible for improved organotypic structure over FA. Though it is unclear, if Matrigel is taken up by the organoid and then embedded as a component in the aggregate or if processes are initiated to increase fibronectin expression. To answer this, we conjugated Matrigel with biotin for visualization of potential incorporation in the organoid (Supplementary Fig. 6a). Cross-sectional confocal images (z-sections) revealed biotinylated Matrigel is indeed embedded throughout the aggregates (Supplementary Fig. 6b-c). Due to a recent report showing that immobilization of Matrigel promotes fibronectin fibrillar formation[54], we immunostained whole FA and hFLO cell aggregates to determine the extent of fibronectin expression and organization into fibrils.

Through semi-quantitative western blot analysis, we observed increased fibronectin expression (Supplementary Fig. 6d). Since there is a known link between fibrillar formation and vascular morphogenesis[55], we used AngioTool[56] (an image analysis software employed to measure network formation) to characterize fibronectin fibrillar morphology and found significantly improved fibrillar features such as branching and lacunarity (Supplementary Fig. 6e-f). Together this data shows that there is both an increase in fibronectin expression and fibril assembly with the supplementation of Matrigel.

### Soluble ECM promotes vascular formation in aggregates

Literature shows that endothelial and fibroblast interactions are essential for vascular-like branching.[47, 48] We used live confocal images to explore the effects of soluble ECM in formation of a vascular-like network. We found that co-culture of epithelial and endothelial cell did not form branching networks but co-culture of epithelial and fibroblast were sufficient for fibroblast network formation (Supplementary Fig. 7a) Presence of all three cell lines with ECM supplementation resulted in the formation of vascular like networks composed of both fibroblast and endothelial cells around epithelial cells (**Fig.1c and 3a**, Supplementary Fig. 2). Quantification using AngioTool verified significantly more vascular like characteristics such as, endothelial-fibroblast branching (**Fig. 3b** and Supplementary Fig. 7b). Thus, for the first time, we show that endothelial branching is activated in the presence of soluble ECM and is dependent on fibroblasts.

**Fig. 3:**
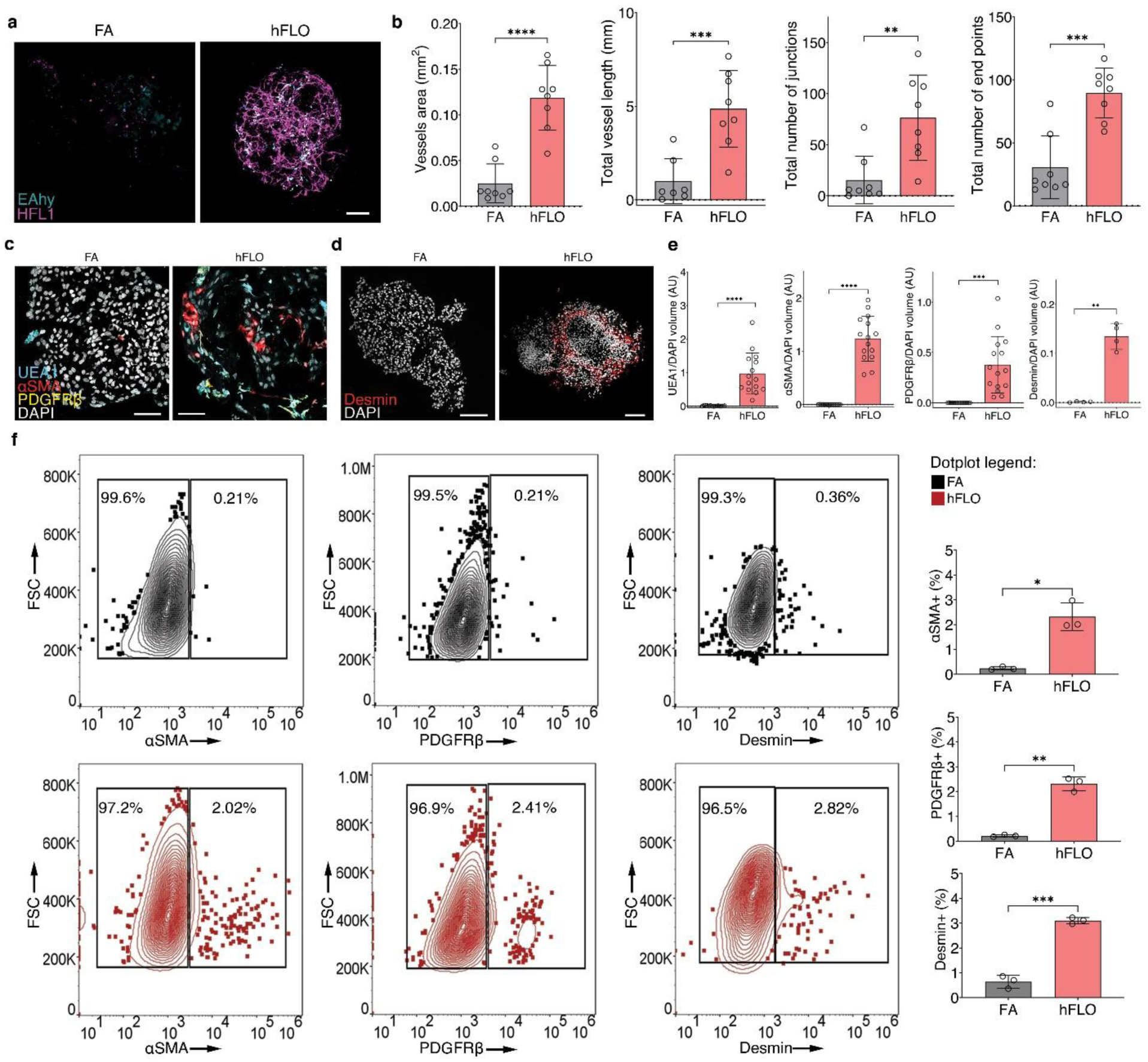
Vascular endothelial branching. **a**, Live fluorescence confocal images (maximum projection) of day 14 cell aggregates showing EAHy (endothelial, blue) and HFL1 (magenta). Scale bars, 100 µm. **b**, AngioTool outputs characterizing the abundance and branching of the EAhy-HFL1 networks. We used 8 independent aggregates per condition. \*\**P*<0.01, \*\*\**P*<0.001, \*\*\*\**P*<0.0001 determined using two-tailed t-test with Welch’s correction. **c**, Fluorescence confocal images (maximum projections) of day 14 hFLO showing the presence of vascular cells including endothelial-like (UEA1 lectin), smooth muscle-like (αSMA), and pericyte-like (PDGFRβ) cells.**d**, Another perivascular marker, desmin was detected in day 14 hFLO. DAPI was used as the nuclear stain. Scale bars, 100 µm. **e**, Volumetric analyses of UEA1, αSMA, PDGFRβ, and desmin in FA and hFLO aggregates normalized to DAPI volume. We used n=15 independent cell aggregates for UEA1, αSMA, and PDGFRβ, and n=4 independent cell aggregates for desmin. \*\**P*<0.01, \*\*\**P*<0.001, \*\*\*\**P*<0.0001 determined using two-tailed t-test with Welch’s correction. **f**, Flow cytometric analysis of whole cell aggregates for expression of αSMA, PDGFRβ, and desmin in FA and hFLO. Contour plots show percent gated cells, contours, and outliers were indicated as dots. Three independent plates were used for each. Bar plots show means, individual points, and standard deviation from positive gates. \**P*<0.05, \*\**P*<0.01, \*\*\**P*<0.001 determined using two-tailed t-test with Welch’s correction.

We used expression and patterning of known markers of vascular maturation (having both endothelial and perivascular cells) to verify formation of vascular like structures. Imaging of vascular cells in hFLO revealed the presence of endothelial cell marker Ulex europaeus lectin (UEA1+), in proximity to perivascular cell markers including smooth muscle actin (αSMA), platelet-derived growth factor receptor (PDGFRβ), and desmin (**Fig. 3c, d**). Upon volumetric analysis, we found that these endothelial and perivascular-like expressing cells were barely present in FA (**Fig. 3e**). Flow cytometry analysis confirmed that although these perivascular-like cells were rare in hFLO, accounting for 2-3% of total cells, they were nearly absent in FA with 0.2-0.36% of the total cells (**Fig 3f)**. This provides additional evidence that growth of organotypic, vascular-like networks are promoted with the addition of soluble ECM.

### Pro-morphogenic expression is enriched with soluble ECM

To determine if the histological and biochemical changes noted thus far are supported by proteomic changes, we analyzed the whole proteome of day 14 FA and hFLO. This data was used to identify upregulated or down-regulated protein groups with addition of soluble ECM (Supplementary Fig. 8a). After imputation and quantile normalization, we confidently identified 967 proteins common to both FA and hFLO (Supplementary Fig. 8b-e). Some of the most abundant proteins detected in hFLO are basement membrane proteins including laminins LAMB1 and LAMC1, which is expected based on the addition of Matrigel (Supplementary Fig. 8e). Upon clustering and principal component analysis we observed statistical difference between FA and hFLO proteome (**Fig. 4a-b** and Supplementary Fig. 8f). Hierarchical clustering of protein-protein correlation across all samples shows the general proteome landscape changes and implies co-regulation among proteins associated with hFLO (Supplementary Fig. 8g). Among the proteins, 147 were determined as differentially expressed proteins but only 42 are highly differentially expressed having an absolute fold change greater than 2 (**Fig. 4c**). 19 proteins (2% of total) were upregulated in hFLO while 23 (2.4% of total) were downregulated. Some upregulated proteins include extracellular structural proteins such as COL18A1, LAMA1, and NID1 supporting the possible deposition of Matrigel in the aggregates as observed in our biotin experiment (**Fig. 4b**). We used the built-in gene ontology term (GO-Term) biological process set in Gene Set Enrichment Analysis (GSEA) to perform an unbiased characterization of the whole proteome (**Fig. 4d**).[57] Some of the top significantly enriched GO-terms in hFLO are pro-morphogenic, related to ECM organization, and markers of active metabolism (**Fig. 4e** and Table S2). In FA, however, the top significantly upregulated GO-Terms are negative regulation of signaling and metabolism (**Fig. 4f** and Table S3).

**Fig. 4:**
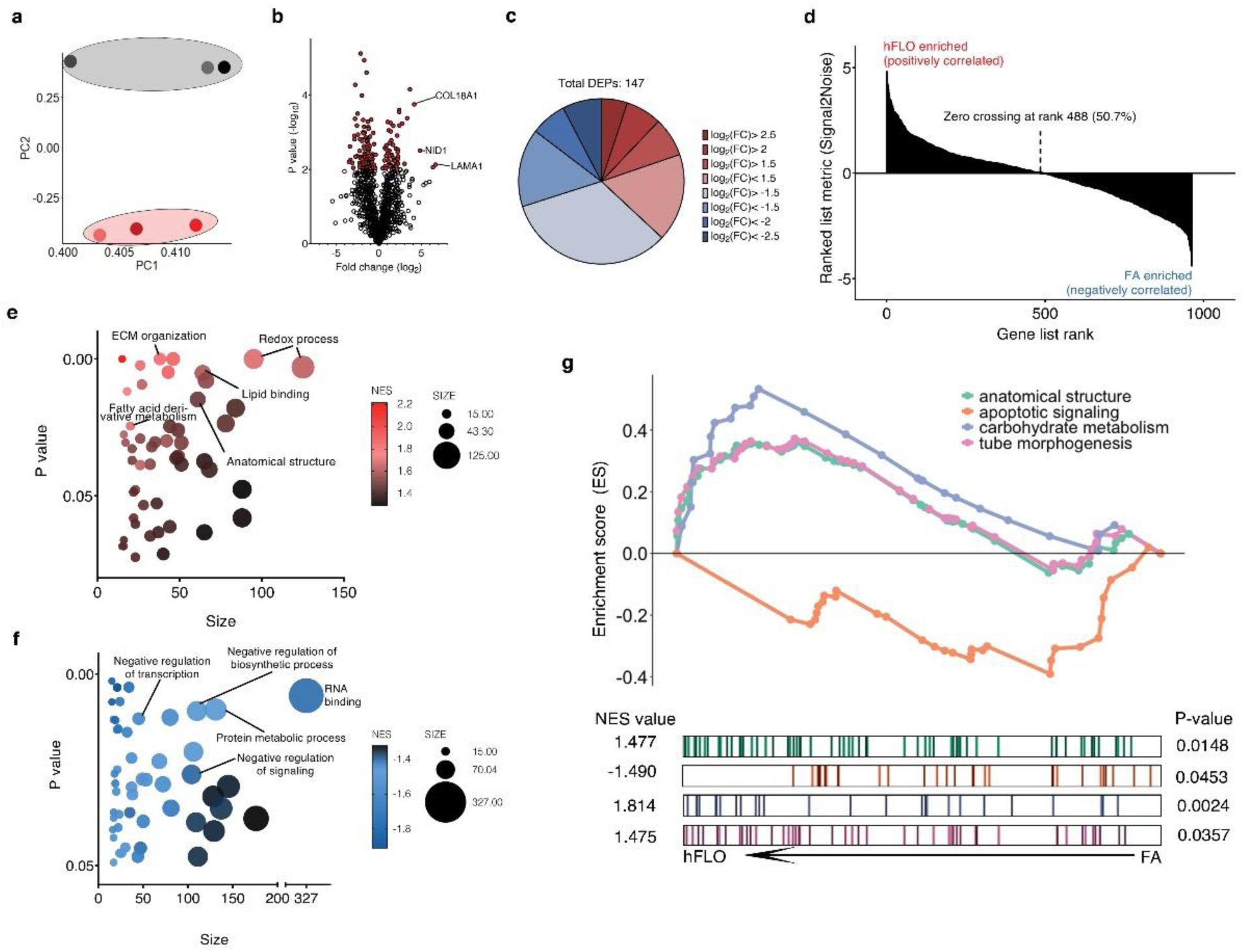
Shotgun proteomics. **a**, Principal component analysis plot of the first and second components. Dots represent the whole proteome of three independent samples used for FA aggregates (grey) and hFLO (pink). Grouping is highlighted using an enclosing oval. **b**, Volcano plot showing fold changes (log2FC) of hFLO versus FA aggregates. *P* values were calculated using two-tailed t-tests with Welch’s correction. Red points indicate proteins with *P*<0.01. **c**, Distribution of the differentially expressed proteins (DEPs, *P*<0.01). **d**, Gene set enrichment analysis (GSEA) plot of all proteins ranked based on enrichment score (Signal2Noise). **e**, Scatter plot of the top 50 GO-Terms enriched in hFLO. Dot size corresponds to number of genes annotated per term and color corresponds to the normalized enrichment score (NES). **f**, Scatter plot of the top 50 GO-Terms enriched in FA (negative enrichment in hFLO). **g**, GSEA enrichment plot for selected GO-Terms. Barcodes show individual proteins annotated in each GO-Term and their respective ranks based on enrichment of hFLO versus FA.

We selected a few relevant up-regulated GO-Terms including anatomical structure formation involved in morphogenesis (GO: GO:0048646, NES= 1.477), cellular carbohydrate metabolism (GO: GO:0005975, NES= 1.814), and tube morphogenesis (GO: GO:0035239, NES= 1.475) for further interrogation of each gene set (**Fig. 4g**). Additionally, we included a downregulated process, intrinsic apoptotic signaling pathway (GO: GO:0097193, NES= -1.490) in this interrogation (**Fig. 4g**). These data support our previous observations that the supplementation of soluble ECM promotes cell survival and the formation of organotypic structures in hFLO triculture aggregates.

### hFLO accurately models pulmonary fibrosis and vascular development

There is a critical need for viable organotypic models for the human lung to model diseases including pulmonary fibrosis (PF). In order to fill this need, we induced fibrosis using bleomycin in our hFLO model. Bleomycin-induced PF is an extensively used animal-based model for fibrosis, despite significant cellular and molecular differences between mouse and human lungs.[58-61] Moreover, therapeutics against PF are limited due to lack of organotypic lung models adapted for drug testing and screening.[60] Translation animal therapeutic research revealed the promise of rho-associated protein kinase (ROCK) inhibition in the reversal of bleomycin-induced lung fibrosis.[61] However, this therapy has not yet been validated in a human lung model. Here, we grew hFLO for 7 days and induced fibrosis using 20 µg mL^-1^ bleomycin for 3d, after which aggregates were treated with 10 µM fasudil, a ROCK inhibitor for 4 days to observe possible changes in fibrosis (**Fig. 5a**). Since we did not include bleomycin from day 10-14 of culture, the effects observed in the bleomycin treatment are most likely the result of the irreversible effects within that timeframe. In PF, there is an observed alveolar and bronchiolar epithelial cell death and fibroblast abundance.[62, 63] We observed that with bleomycin induction, there was a decrease in the epithelial A549 with an increase in fibroblast HFL1 population (Supplementary Fig. 9b). Moreover, we observed decreased lumen area in bleomycin-treated aggregates which is rescued by the fasudil treatment (Supplementary Fig. 9c).

**Fig. 5:**
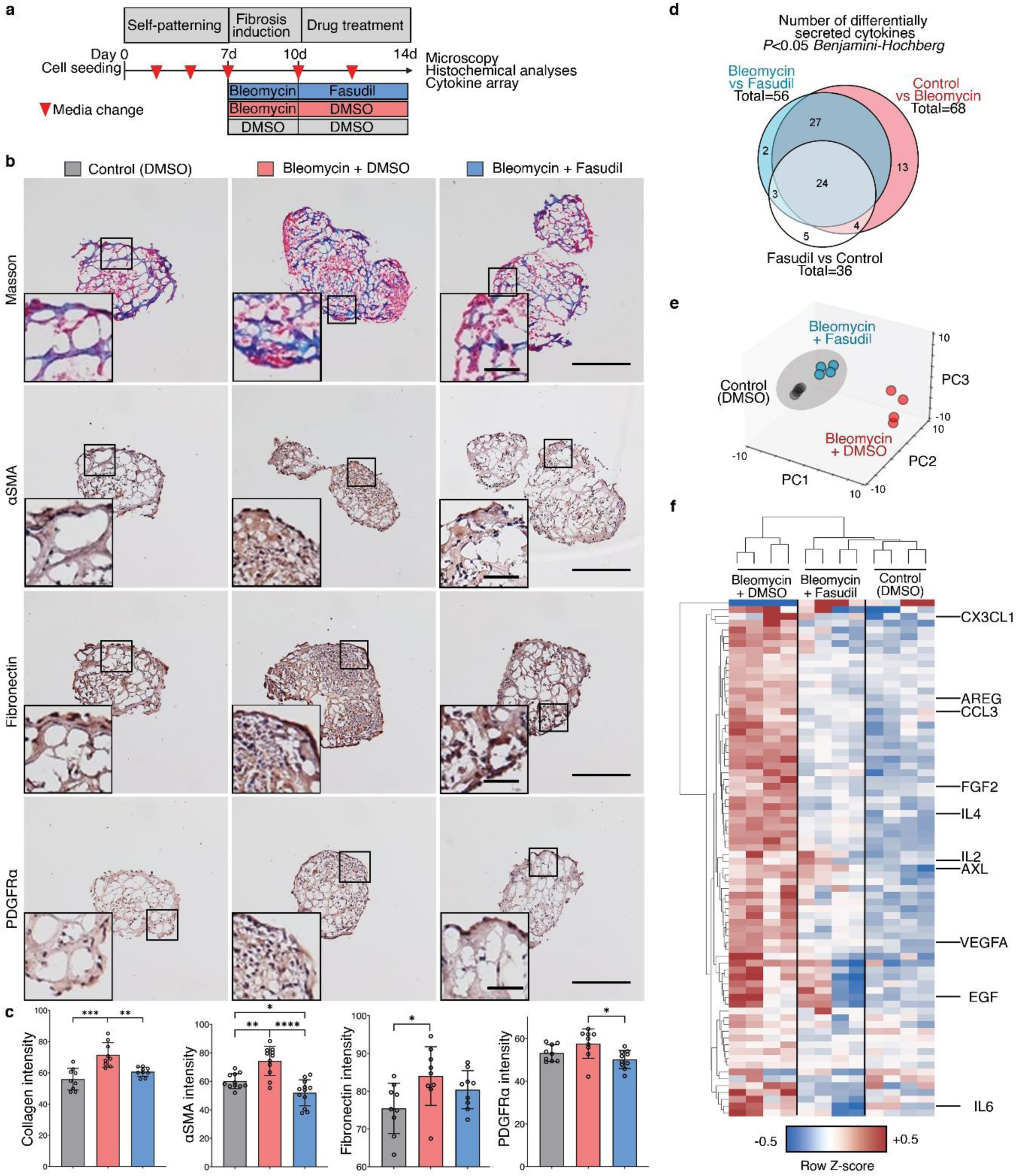
Pre-clinical hFLO-bleomycin fibrotic model and resolution of fibrotic features using fasudil. **a**, Schematic diagram of the workflow showing aggregate formation and self-patterning for 7 days, induction of fibrosis using bleomycin (20 µg/mL) for 3 days, and antifibrotic trial using fasudil (10µM). Dimethyl sulfoxide (DMSO) is the vehicle used for the bleomycin and fasudil. **b**, Histochemical and immunohistochemical analyses for the extent of induced fibrosis in the hFLO aggregates. Insets show fibroblastic foci for bleomycin-treated aggregates. Scale bars, 300 µm for full image; 60 µm for inset. Masson staining shows the extent of collagen deposition (blue). We used 9 independent aggregates per condition. Hematoxylin and DAB immunostaining (H-DAB) of pro-fibrotic markers αSMA, fibronectin, and PDGFRα, respectively. We used n=11, n=9, and n=9 independent aggregates per condition for H-DAB probing for αSMA, fibronectin, and PDGFRα, respectively. **c**, Histochemical and immunohistochemical image analyses. Bar graphs show means, individual values, and standard deviations are shown. \**P*<0.05, \*\**P*<0.01, \*\*\**P*<0.001, \*\*\*\**P*<0.0001 determined using One-way ANOVA with Welch’s correction. Color legends shown in **b. d-f**, Cytokine array profiling the secreted factors from the treatment groups. Raw immunoblot intensity were normalized to internal positive and negative controls. **d**, Differentially secreted factors (*P*<0.05) were determined using two-tailed Welch’s t-test with Benjamini-Hochberg FDR correction. Statistical tests were done comparing each group to each other. Venn diagram shows overlap among differentially secreted factors. **e**, Principal component analysis of the differentially secreted factors. Dots show media samples collected from the organoids and axes represent principal components (PC) 1, 2, and 3. Clustering among fasudil-treated and control samples is emphasized by the gray ellipse. **f**, Cluster heatmap (Euclidean) of the differentially secreted factors showing clustering of fasudil-treated and control samples, separate from bleomycin-treated samples. Known cytokine/chemokine effectors of angiogenesis and fibrosis are annotated. Cell color is based on row average-normalized z-scores.

We used immunohistochemistry to ascertain whether fasudil treatment leads to decreased expression of common fibrotic markers, including collagen deposition, expression of stress fiber marker αSMA, ECM marker fibronectin, and fibroblast marker PDGFRα. We observed and quantified decreased intensity of collagen deposition using Masson staining in organoids treated with fasudil (**Fig. 5b** and Supplementary Figure 9d). Moreover, the increased pro-fibrotic expression after bleomycin insult was lowered after fasudil treatment (**Fig. 5c-e** and Supplementary Fig. 9d). Localization of these markers also led to the identification of fibroblastic-myofibroblastic foci in the aggregates, another histopathological feature of IPF (**Fig. 5c-e**).[64]

To further confirm the fibrosis in hFLO-bleomycin fibrotic model and the anti-fibrotic effects of fasudil, we collected culture media from the aggregates and performed a cytokine array using 120 common inflammatory factors. Among these, we found that 68 factors were significantly increased by the bleomycin treatment (Control vs Bleomycin, **Fig. 5d** & Table S6). While 56 factors were significantly changed associated with the fasudil treatment (Fasudil vs Bleomycin) and 36 factors between control and fasudil treatment (**Fig. 5d**). Among these significantly secreted factors, 51 are common between the factors enriched by bleomycin insult and the factors that are changed by fasudil treatment. Upon data reduction using principal component analysis, we observed a closer clustering of the fasudil-treated fibrotic aggregates with the control (vehicle, DMSO-treated) aggregates but not with the bleomycin-insulted aggregates (**Fig. 5e**). This is further supported by hierarchical clustering showing the association of the fasudil-treated aggregates with the normal, non-fibrotic aggregates in a single clade, while the bleomycin-treated hFLOs are grouped separately (**Fig. 5f**). This shows very similar patterns of cytokine secretion were observed in the control and fasudil-treated fibrotic aggregates. Upon close examination of these secreted factors, we observed that pro-inflammatory and pro-angiogenic factors are increased in bleomycin treatment. Fasudil treatment significantly decreased these pro-inflammatory factors associated with tissue fibrosis including: CX3CL1, AREG, CCL3, IL4, IL2, AXL, EGF, and IL6 among others (**Fig. 5f**). Moreover, we observed increased secretion of pro-angiogenic factors including FGF2 and VEGFA in bleomycin-insulted aggregates which is then decreased with fasudil treatment. These results suggest that hFLO can accurately model certain fibrotic features *in vitro* and potentially evaluate pre-clinical anti-fibrotic drugs, as illustrated here with fasudil.

We then looked into another application of hFLO, a model for hypoxia-induced angiogenesis[65] which can mature to full perfusability[66]. Blood vessel formation is an important process which affects several lung diseases including cancer[67] and pulmonary hypertension[68]. Previous vascular organoid methods use of hypoxia (5% O_2_) and supplementation of proangiogenic factors vascular endothelial growth factor (VEGF) and fibroblasts growth factor 2 (FGF2) to induce vascular endothelial branching, differentiation and maturation.[69] Thus, at day 7, hFLO was treated with 60ng mL^-1^ FGF2 and 60ng mL^-1^ VEGF and cultured in 5% O_2_ (**Figure 6a**).

**Fig. 6:**
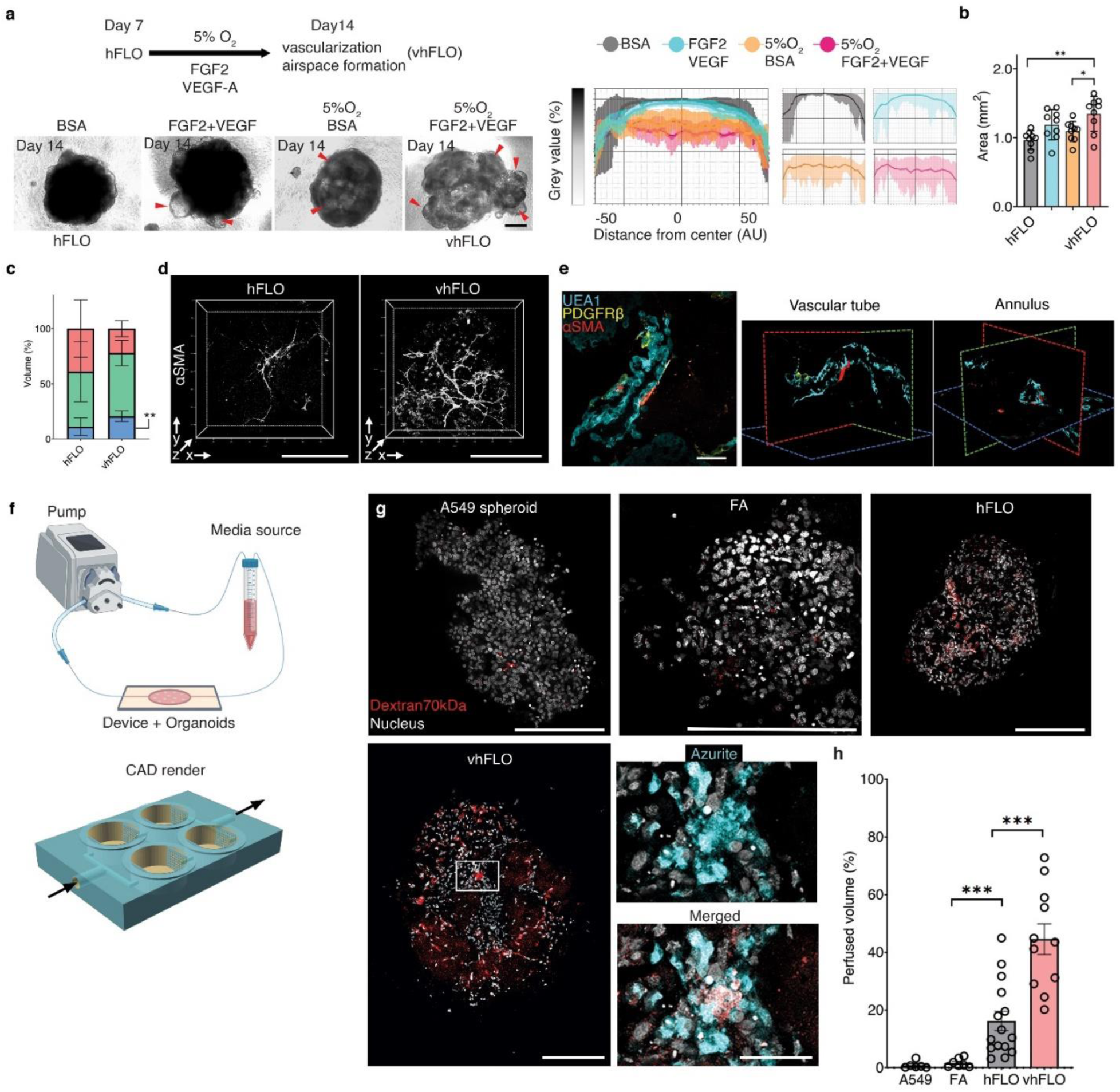
Hypoxic vascularization using hFLO. **a**, Schematic diagram of the workflow showing further vascular growth using hypoxia and pro-angiogenic factors (top left). Bright-field images of aggregates at day 14 of culture (bottom left). vhFLO is hFLO grown under hypoxia for 7 days with supplementation of FGF2 and VEGF. Line profile analysis at the major axis of the aggregates expressed as normalized grey value. Effects of the hypoxic culture in aggregate density is shown using line profile analysis (right). Lines indicate the grey value range at the specific normalized distance from the center. Scale bar, 300 µm. **b**, Aggregate area at day 14 of culture. 9 independent aggregates were measured for each condition. \**P*<0.05, \*\**P*<0.01 determined using One-way ANOVA with Welch’s t-test. **c**, HFL1 (red), A549 (green), and EAhy (blue) cell population percent volume composition in hFLO and vhFLO. N=7 independent 3D live confocal images were used. \*\**P*<0.01 determined using two-tailed t-test with Welch’s correction. **d**, Whole aggregate immunostaining of αSMA expression. Scale bar, 200 µm. **e**, 3D fluorescence confocal image of vhFLO showing vascular-like formation and localization of endothelial-like (UEA1 lectin+, BLUE), pericyte-like (PDGFRβ+, yellow), and smooth muscle-like (αSMA+, red). 3D renderings highlight annular tube morphology typical of a vasculature. Scale bar, 50 µm. **f**, Illustration of the vascular perfusion assay using dextran 70 kDa flown through a peristaltic pump system (top). The chip used for this assay is shown as a CAD render with fluidic flow indicated by arrows. Additional information could be found in Supplementary Fig. 10. **g**, Whole aggregate fluorescence confocal images (single z section) of A549 spheroid, FA, hFLO and vhFLO. Images show extent of dextran 70 kDa perfusion. Zoomed images showing dextran and azurite colocalization in vhFLO. Scale bars, 200 µm for full images; 50 µm for zoomed images. **h**, Volumetric analysis of extent of perfusion in the aggregates. A549 spheroid was used as a negative control. We used n=6 A549 spheroid, n=7 FA, n=15 hFLO, and n=11 vhFLO independent aggregates. ***p<0.001 determined using One-way ANOVA with Welch’s t-test.

Hereafter, we will refer to the hFLO treated with FGF2 and VEGF under hypoxia as the vascular hFLO (vhFLO). In vhFLO, we observed hallmarks of lung development such as the increased prevalence of lumen and increased aggregate size with FGF2 and VEGF treatment under hypoxia when compared to hFLO (**Fig.6a-b**). We also observed an increase in EAhy population volume and junction formation in vhFLO (**Fig. 6c** and Supplementary Fig. 10). We then tested whether this increase in EAhy resulted in improved vascular branching and maturation. Immunostaining revealed increased prevalence of αSMA+ networks (**Fig. 6d**). 3D confocal imaging revealed luminal tube-like formation of UEA1+ endothelial cells with pericyte-like (PDGFRβ+) and smooth muscle-like (αSMA+) cells surrounding the abluminal side (**Fig. 6e**).

An important feature of a mature vasculature is perfusability.[70] We created a PEGDA biocompatible 3D-printed chip to assess vascular perfusability using 70kDa Dextran (**Fig.6f** and Supplementary Fig.11).[35] Upon fluidic exposure to fluorescent 70kDa Dextran, we observed that hFLO and vhFLO had perfusable regions while A549 and FA spheroids did not (**Fig. 6g-h**). Among these, vhFLO has the highest perfusable volume, with a 27-fold and 2.7-fold increase from FA and hFLO, respectively. In vhFLO and hFLO, the perfusion was localized with EAhy cells suggesting that perfusion is targeted to vascular-like regions (**Fig. 6g**). These data show that vasculature in hFLO is perfusable and exposure to hypoxia and pro-angiogenic growth factors increase the perfusability of the aggregates, as observed in vhFLO. These findings open additional avenues for the adaptation of hFLO in vascular modeling and angiogenic diseases such as pulmonary hypertension/vasculopathy and in lung cancer.

## Discussion

The role of the ECM in organotypic structures has been extensively studied but research is limited when it comes to the roles and use of soluble ECM. Previous organ model research focused on the constructive (adhesion and architectural biomaterial) and instructive (cell signaling) roles of the ECM as effect of its solid hydrogel form.[14] Regarding its instructive roles, the ECM is a known agonist to various receptors which activate mechanotransduction pathways responsible for organotypic growth, survival, and development.[71, 72] We questioned whether these known organotypic effects of the ECM are true even in its soluble form. To study this possibility, we developed a novel hybrid suspension culture that features organotypic growth instead of the typical compacted spheroid morphology with the help of soluble ECM. To show the efficacy of this novel 3D method, we introduced soluble ECM (growth factor reduced Matrigel) to a triculture of epithelial, endothelial, fibroblast cells. With this new methodology, an organotypic lung vascular alveolar model was derived using stable cell lines. After 14 days in our soluble ECM triculture, we observed various cell populations including type I-like (Pdpn+), type II-like (SPC+, lamellar body +) alveolar epithelial cells, mesenchymal cell populations including fibroblasts, endothelial cells, pericyte-like (PDGFRβ+), and smooth muscle-like cells (αSMA+, desmin+) cells. Moreover, the distribution of these cells was similar to that of the alveolus and the blood vessels. Early studies and recent advancements in single cell RNA sequencing show that these cell types populate the lung alveolus and the associated mesenchyme and vasculature.[26, 73] Other novelties of this research include: 1) formation of organotypic luminal structures in a suspension culture where normally, cells form compact tissue-like spheroids; 2) effects of soluble ECM in 3D growth, cell survival, and long-term structure maintenance; 3) the first perfusable vasculature in an organotypic lung model; and 4) creation of a high-throughput bleomycin lung alveolus model along with human cell-based pre-clinical study of an anti-fibrotic drug.

Recently, a study illustrated the use of a soluble ECM, tropoelastin in promoting growth and survival of mesenchymal stem cells in 2D culture.[11] Another study showed that soluble collagen VI is able to prevent apoptosis in serum-starved fibroblasts.[74] However, these pro-survival effects of soluble ECM have not been demonstrated in 3D culture. Within the field of 3D culture overcoming cell death is vital to create viable cell aggregates capable of sustaining long term growth. Here, we present evidence that soluble ECM improves cell survival in 3D suspension culture. During the 14-day culture, proteomic data gives insights on the feasible pro-survival mechanism of the soluble ECM to be hinged on increased carbohydrate metabolism[75] and the prevention of anoikis[76], two processes that are well-studied effects of solid ECM. Our growth assays suggest that culture in soluble ECM allows for long-term survival and structural maintenance of cell aggregates up to 70 days of culture. The ability of 3D models to be maintained for an extended period of time is essential in studying long-term processes such as organ maturation, disease resolution, and aging among others. We believe that our finding is the first account to show the pro-survival effects of non-solid ECM in 3D multicellular aggregates. One of the major feature that sets organoids apart from spheroids is the presence of central lumen which is essential in organs for their specialized functions.[77] In the lung, the alveolar lumen is populated by type I and type II epithelial cells which facilitate gas exchange and pressure regulation, respectively.[78] Moreover, lumen formation is not just confined to the epithelium but is also observed in the vasculature which allows for efficient material circulation.[79, 80] In the hFLO, we observed both types of lumina formed via immunochemistry, histology, and electron microscopy. In the epithelial lumen, we observed the presence of multilamellar body-presenting type II-like cells in close proximity to long, slender type I-like cells. Moreover, upon close examination, we also observed that adjacent to these epithelial cells and separated by a basement membrane are luminal endothelial cells (LECs). Further, immunostaining and flow cytometry revealed that at least three types of cells inhabit these lumina—endothelial cells, pericytes, and vascular smooth muscles.

Current stable cell suspension co-cultures lack the presence of perfusable and branching vasculature, thus, these cell culture methods are only capable of recapitulating compacted organization and are growth-limited due to poor material circulation.[77, 81-83] In the presence of solid ECM endothelial cells alone form networks and co-culture with fibroblast cells result in lumen formation of endothelial cells.[47, 48] In soluble ECM culture, we show that endothelial cells only form networks in the presence of fibroblasts. Additionally, these endothelial-fibroblastic branching networks express perivascular cell markers. This observed vascularization followed by maturation into functioning perfusable tubular networks is vital for material circulation and for epithelial-endothelial interaction to form. During endothelial tube formation, fibronectin (FN) is required, and its organization into fibrils initiates vascular tube formation.[55]. Lu *et al*. show that Matrigel promotes FN fibrillogenesis only when it is immobilized and not in its soluble form.[54] In our model, we observed FN fibrillogenesis even with the use of soluble Matrigel due to the immobilization and incorporation of the soluble ECM into the aggregate. This immobilization magnifies the assembly of fibronectin into fibrillar structures, alluding to the instructive role that the immobilized ECM has in fibril assembly.[54] These vascular interactions are the basis of the alveolar gas exchange unit, thus the importance of these results become clear.

Recent developments in tissue engineering led to the creation of induced pluripotent stem cell (iPSC)-derived lung organoids.[84] These organoids show high-fidelity in mimicking organ development, and have a high potential for use in disease modeling and subsequent drug development.[85] Moreover, these iPSC-derived lung organoids show high organotypic complexity with multiple types of cells from various lung regions including the bronchi and the alveolus.[86, 87] Though, directed differentiation of iPSC-derived organoids is a laborious process which takes on average over 50 days, involves several culture transfers, and requires multiple growth factors.[88] In contrast, hFLO features a minimalistic methodology and a 14-day culture time to produce aggregates with organotypic lung alveolar markers. This characteristic makes this model adaptable for automated downstream high throughput studies. These features also make hFLO a feasible and important tool for time-sensitive disease modeling like the COVID-19 pandemic or future respiratory pandemics. Moreover, the lack of a perfusable vasculature in iPSC derived organoids limit their expansion and their use in vascular lung disease research. We posit that the use of the basic vascular unit described in this paper is vital in the improvement of current epithelial organoids in expanding their growth and stability via co-culture.[89]

To illustrate a disease modeling application of hFLO, we used bleomycin induction to model lung fibrosis. Bleomycin promotes inflammation and formation of fibrotic lesions with similar biochemical and morphological features in patients with IPF and other fibrotic lung disorders.[90] Histopathological and biochemical patterns of usual interstitial pneumonia (UIP), the hallmark feature of IPF include: epithelial injury leading to disrupted basement membrane (ie. increased collagen deposition), loss of airspaces, presence of dominant fibroblastic foci, and increased mesenchymal markers.[59, 91] We found that bleomycin induction in hFLO resulted in increased collagen deposition implying disruption in the basement membrane. The bleomycin-treated hFLO organoids had decreased air space-like gas exchange units as shown through lumen analysis, presence of fibrotic foci and increased mesenchymal markers. Moreover, the aberrant cytokine secretion in the hFLO-bleomycin is enriched with pro-inflammatory cytokines implicated in fibrosis, especially in PF; these includes CX3CL1[92], AREG[93, 94], CCL3[95, 96], IL4[97], IL2[98], AXL[99, 100], EGF[101], and IL6[102] among others. Our observations demonstrate that the hFLO-bleomycin fibrotic model not just recapitulates alveolar histopathological features of PF, but also the associated pro-fibrotic, inflammatory cytokine secretion. Together, these results suggest that the bleomycin induction in hFLO successfully achieved fibrosis and is irreversible during the timeframe but was ameliorated by fasudil treatment. The application of this innovative method closely replicates multiple features in patients with PF and since hFLO is a human *in vitro* model, it has the potential to replicate disease pathogenesis and drug interactions that current animal models are unable to show.

We chose to test our human lung fibrosis model with fasudil, a ROCK inhibitor clinically used to treat cerebral vasospasm in Japan and China, but not yet approved in the United States.[103, 104] Fasudil was found effective in resolving PF symptoms in traditional 2D *in vitro* and *in vivo* mouse studies. However, lack of data in a human model of the disease precludes further investigation and fasudil’s potential adoption as a human PF treatment.[105, 106] Our results show that fasudil improved the above-mentioned PF pathologies present in the hFLO-bleomycin model. Moreover, the highly similar cytokine secretion of fasudil-treated fibrotic organoids to non-disease hFLO indicate the reversal of the inflammatory fibrotic effects of bleomycin. These results may also open the door to pre-clinical trials of existing ROCK inhibitors such as Y27632 and the development of other ROCK inhibitors. Further isoform-specific ROCK inhibition studies are needed for more fine-tuned targeting of the disease.[107] We view the hFLO-bleomycin model to be an important tool that could be used in high-throughput drug screens for PF and related diseases.

To further improve upon the hFLO, we mimicked developmental lung angiogenesis through a combination of pro-vascular growth factors (VEGF, FGF2) and a hypoxic environment (5% O_2_), creating vhFLO, which has more perfusable volume than hFLO. This adds additional physiologically relevant characteristics for researchers to utilize in the study of alveoli-vascular interactions. Our research demonstrates that soluble ECM suspension promotes the formation of a viable vascular lung organoid; further research needs to be completed to further understand the capabilities and limitations of this method. We view that the following steps are necessary to better understand the capabilities of the system: (1) the utilization of a precise serum free, chemically defined media to develop the organoids; (2) the utilization of primary alveolar epithelial cells or adult lung progenitor and stem cells; (3) profiling of all the cell types in the hFLO via single cell RNA sequencing; and (4) formation of macroscopic tissue constructs using hFLO as the building block. Nevertheless, we have illustrated several novelties of our methodology leading to the creation of a human PF bleomycin model, which shows a closer approximation to distal lung than has been shown to date for PF *in vitro* modeling.

## Conclusions

Our hybrid soluble ECM-based 3D culture method promotes organotypic growth using stable mature cells within 14 days of culture. We illustrate that soluble ECM can be used to enhancing 3D growth and survival in suspension cultures. We believe that this is the first account illustrating that soluble ECM can create an organotypic lung model from mature stable cells. Moreover, the lung model presented in this paper has a perfusable vasculature which current lung organoids lack. We demonstrate the utility of the novel hFLO model provide further insights in the pathogenesis of PF. The organotypic features of this model allowed us to mimic PF and demonstrate its resolution using a ROCK inhibitor, fasudil. The application of this method in the formation of patient- or primary-derived organoids is yet to be performed but will be necessary to create high-throughput screens for personalized medicine. Our findings open an avenue for novel tissue engineering using soluble ECM and the application of the hFLO in modeling lung diseases such as PF and COVID-19.

## Acknowledgments

The Simmons Center for Cancer Research Fellowship to J.C.V; and the Earl M. Woolley Research Innovation Award to P.M.V.R. G.P.N. and K.A.C. acknowledge support from the National Institutes of Health (R15GM123405-02) for 3D printer, resin, and device development. Circle V Meats Co. (Springville, Utah, USA) for providing the porcine lungs. Cecilia Sanders for help in editing the manuscript, Mawla Boaks for help in perfusion device 3D printing, and Michael Standing for help in electron microscopy prep and imaging.

## Author Contributions

J.C.V., G.R., and P.M.V.R. conceived, supervised, and acquired funding and resources for the project. Project methodologies were developed by J.C.V.; Cell culture optimizations performed by J.C.V., and C.J.K.; Cell culture-based experiments for the paper were performed by J.C.V., N.A.F. C.G.C., D.A.J., E.L.D., C.J.K., B.M.H., M.S., and J.A.S.; P.D.P., B.M.H., N.A.F., J.C.V. and J.A.S. performed the mouse and pig organ experiments; J.C.V. performed the confocal microscopy imaging; B.M.H., E.L.D., C.G.C., J.C.V., and J.A.S. performed the immuno- and histochemistry; S.R.G. and J.C.V. prepared the samples for and performed the electron microscopy; J.C.V., M.L.V.Z., and C.J.K performed the western blots; J.C.V., B.M.H, N.A.F., C.G.C., and S.R.G. performed the growth assays; J.C.V., C.J.K., C.G.C., and M.T.S. performed the proteomic experiments, visualizations, and bioinformatic data analyses; J.C.V., B.C.J., D.J.J., C.G.C., and B.C. performed the fluidic perfusion experiment. K.A.C. and G.P.N. designed and printed the microfluidic chips; Visualizations and graphs were created by J.C.V., M.T.S., C.G.C., P.D.P., D.A.J., and B.C.J.; Data were curated by J.C.V., N.A.F. C.G.C., D.A.J., E.L.D., C.J.K. S.R.G., B.C.J., B.M.H., P.D.P. and M.T.S.; B.C.K., J.C.V., and A.S. performed the cytokine array and analyses; J.C.V. wrote the initial draft. J.C.V., P.M.V.R., E.L.D., P.D.P., N.A.F., D.J.J., K.A.C., G.P.N., A.S.N., and G.R. reviewed the manuscript and edited the draft; J.C.V., P.V.R., A.S.N., and G.R. revised the manuscript. All authors have read and approved the manuscript.

