## Supplementary for "Soluble ECM promotes organotypic formation in lung alveolar model"

SUPPLEMENTARY FIGURES:

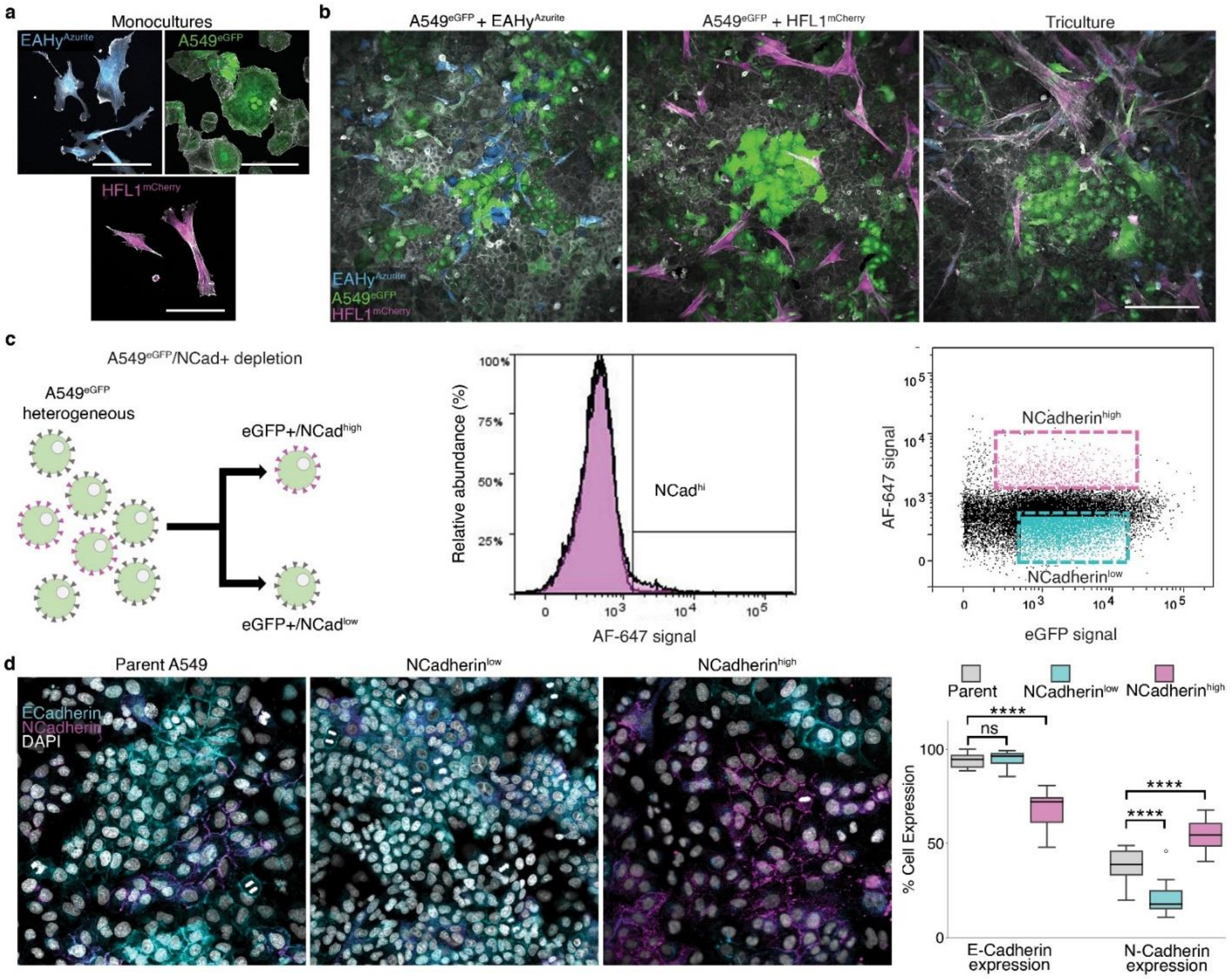


**Supplementary Fig. 1: Cell lines and depletion of NCad high population in A549.**

**a**, 2D monocultures of $EAhy^{Azurite}$, $A549^{eGFP}$, and $HFL1^{mCherry}$ cells. Scale bars, 100 µm. **b**, 2D bicultures of A549 with EAhy (Left) or HFL1 (Middle); and 2D triculture of the cells (Right). Scale bar, 200 µm. **c**, FACS depletion of NCadherin+ cells in A549-eGFP to improve epithelial phenotype. Workflow for the purification (Left). NCad-AF647 intensity histogram (Middle) indicating $NCad^{high}$ population as gated. Delineating NCadherin high and low expressing populations (Right). **d**, These cells were then cultured and after 4 passages stained to confirm phenotypic stability from sorting. ECadherin+(CDH1) and NCadherin+ (CDH2) cells were counted from 15 fields of view and expressed as percent in a box plot.

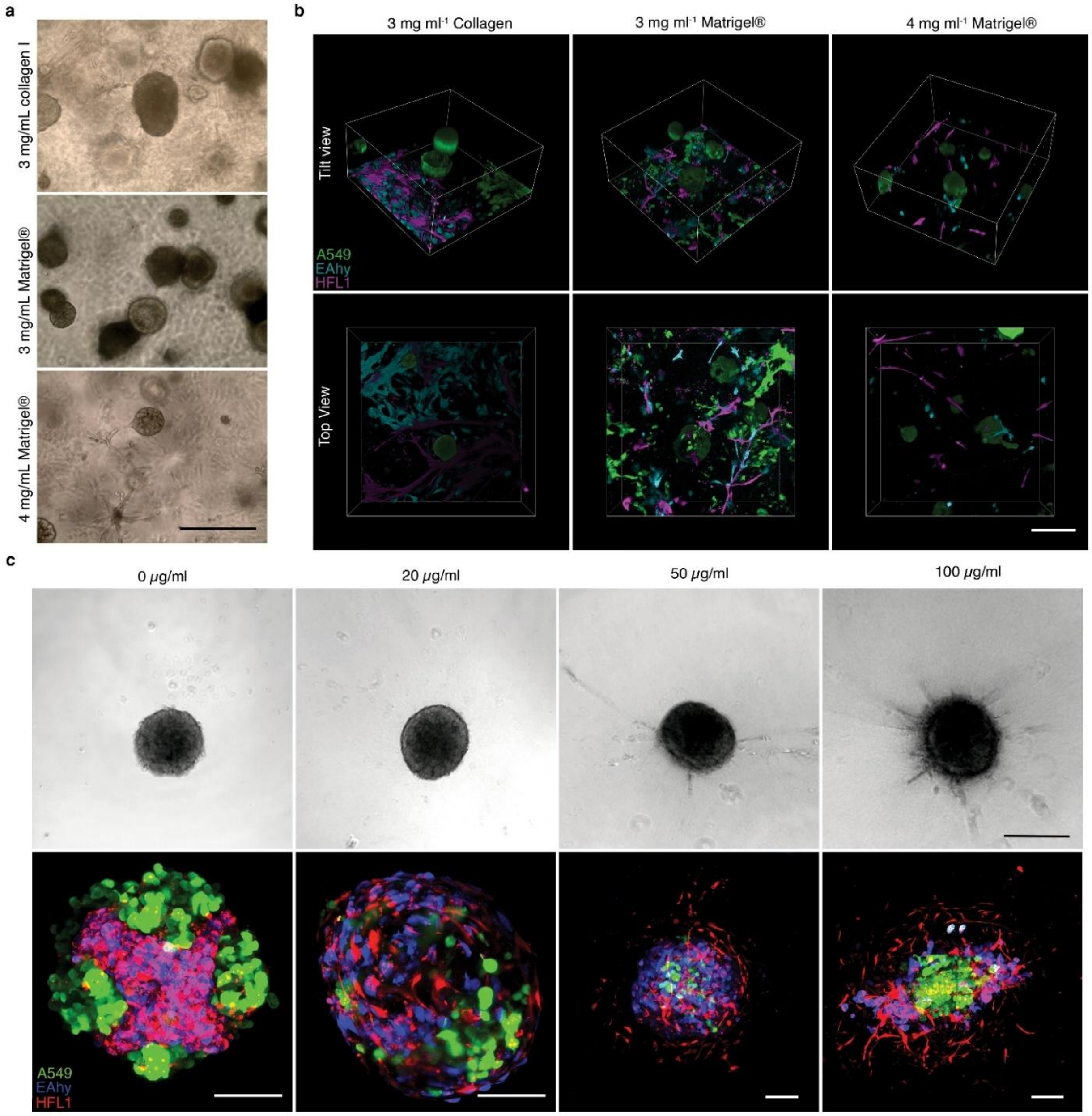


**Supplementary Fig. 2: Effects of ECM concentration in self patterning.**
**a**, Bright field images of tricultures grown embedded in gel ECM: 3 mg ml$^{-1}$ collagen type I, 3 mg ml$^{-1}$ Matrigel, and 4 mg ml$^{-1}$ Matrigel. Scale bars, 300 µm **b**, Live 3D fluorescence confocal images (3D renders, tilted and top view) of the gel ECM cultures. Scale bars, 200 µm. **c**, Effect of collagen type I concentration in self-patterning. Vascular (endothelial-fibroblast) networks sprout from the compact core as shown in the bright field images (Top). Scale bars, 200 µm. Live 3D confocal images (maximum projection) of the cell aggregates. Scale bars, 100 µm.

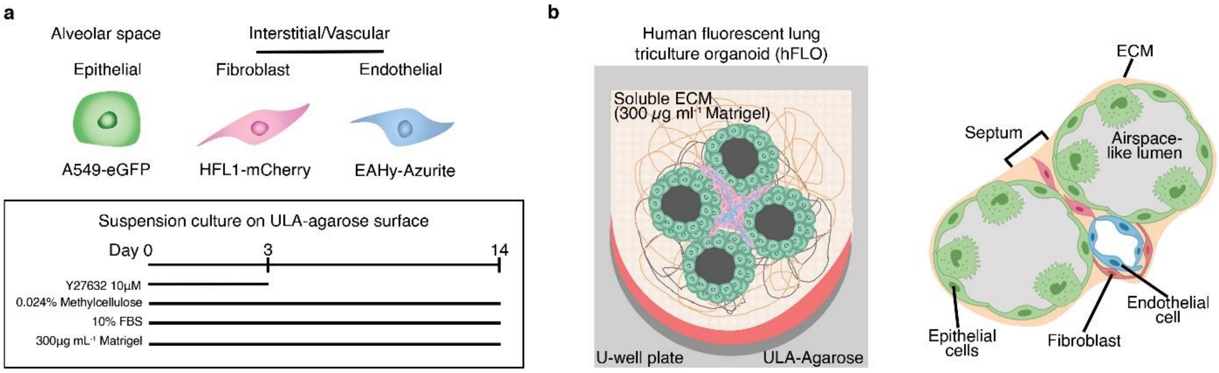


**Supplementary Fig. 3: Suspension culture for a lung organoid via triculture.**
**a**, Cell lines used in the 3D model: epithelial (A549-eGFP) and interstitial/vascular (HFL1-mCherry and EAhy-Azurite). Timeline and culture workflow for culture with 300 µg $mL^{-1}$ Matrigel and 0.024% methylcellulose (Bottom). **b**, A cartoon of the human fluorescent lung triculture organoid (hFLO) depicting suspension culture on ultra-low adherence (ULA) agarose U-bottom well (Left). Microscopic features of the hFLO showing epithelial cells forming septate airspace-like lumen, endothelial cells and fibroblasts interacting with each other in the interstitial spaces, and presence of ECM proteins in the organoids (Right).

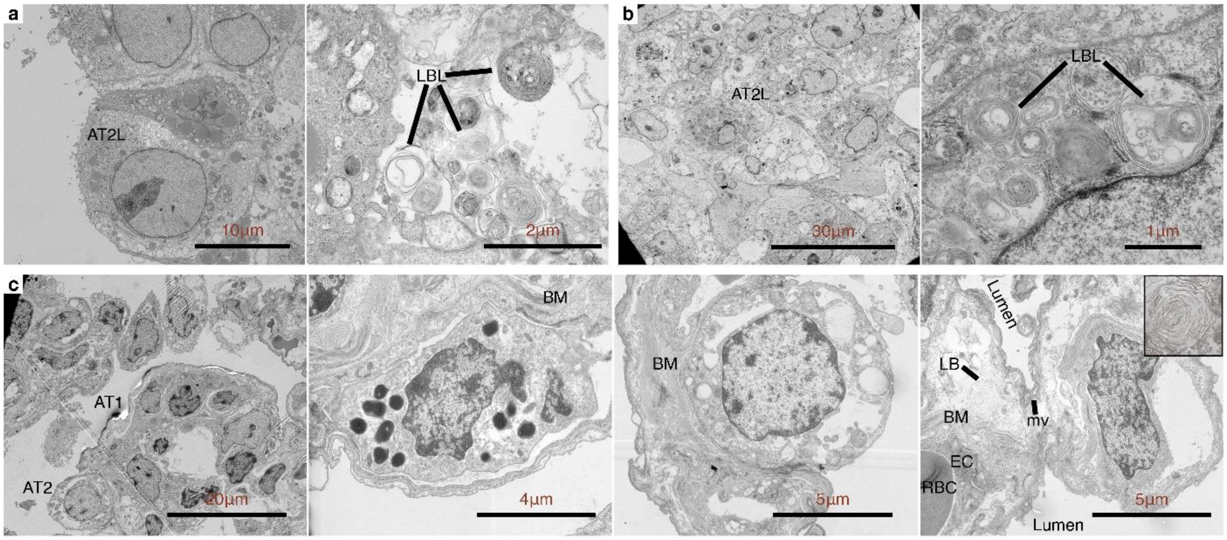


**Supplementary Fig. 4: Transmission electron microscopy.**

**a**, 7-day A549 spheroid cells showing the presence of lamellar body-like inclusions (LBL) typical of a type II-like alveolar cell (AT2L). **b**, 14-day FA aggregates also shown with the presence of LBL. **c**, Porcine lung alveolus showing AT2 cells and AT1 cytoplasmic extension and other features including lumina, basement membranes, EC, red blood cell (RBC), and lamellar bodies (LB). Scale bars as indicated in each pictograph.

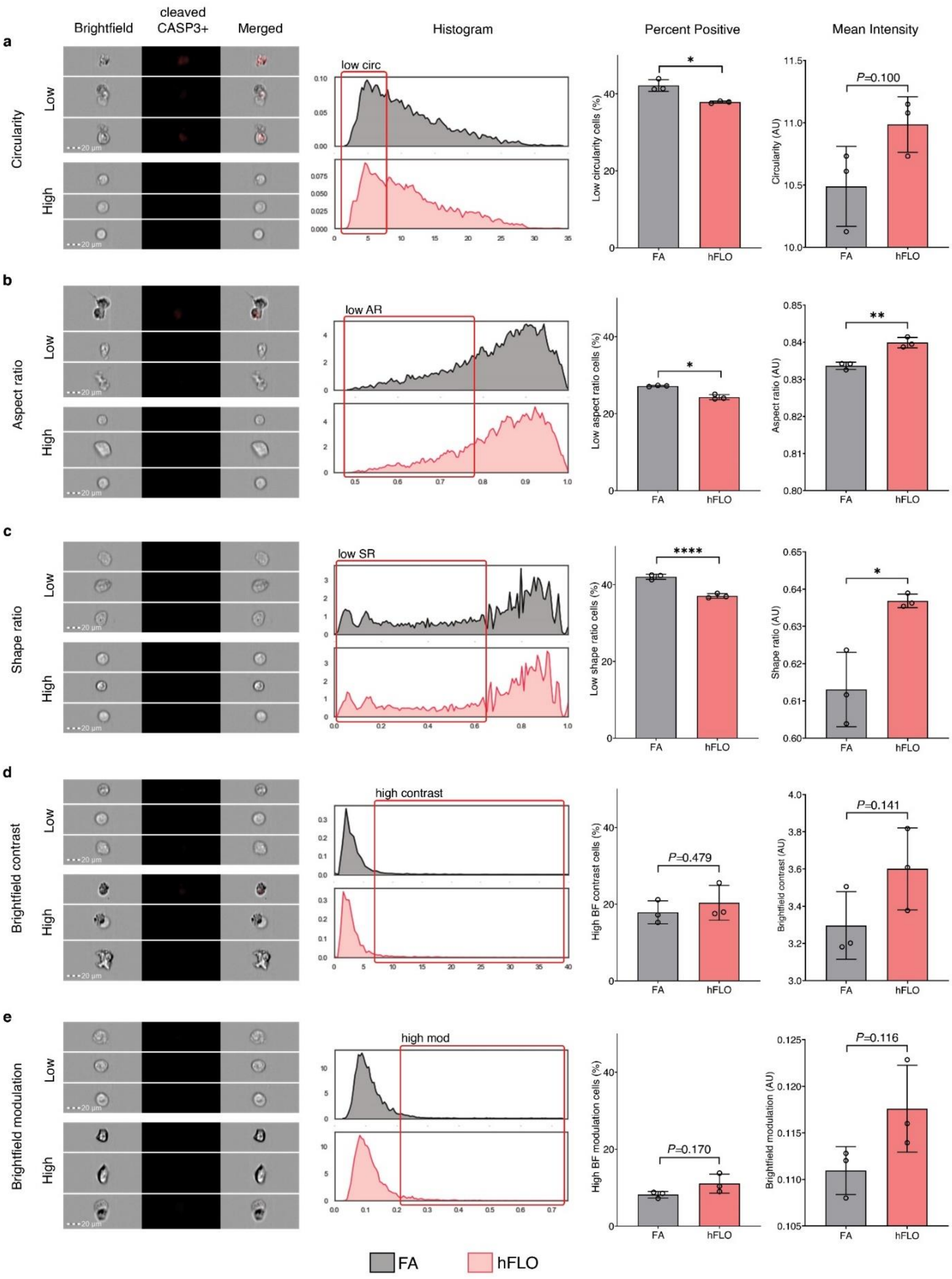


**Supplementary Fig. 5: Morphometric analysis of cell death using imaging flow cytometry.**

**a-e**, Brightfield images representing spheroids typical of high and low spectrums of respective morphological features. Histograms show distributions of respective features of a typical sample of FA and hFLO. Barplots display histogram statistics (left column) and means of feature values (right column). *$p<0.05$; **$p<0.01$; ****$p<0.0001$; Welch's t-test.

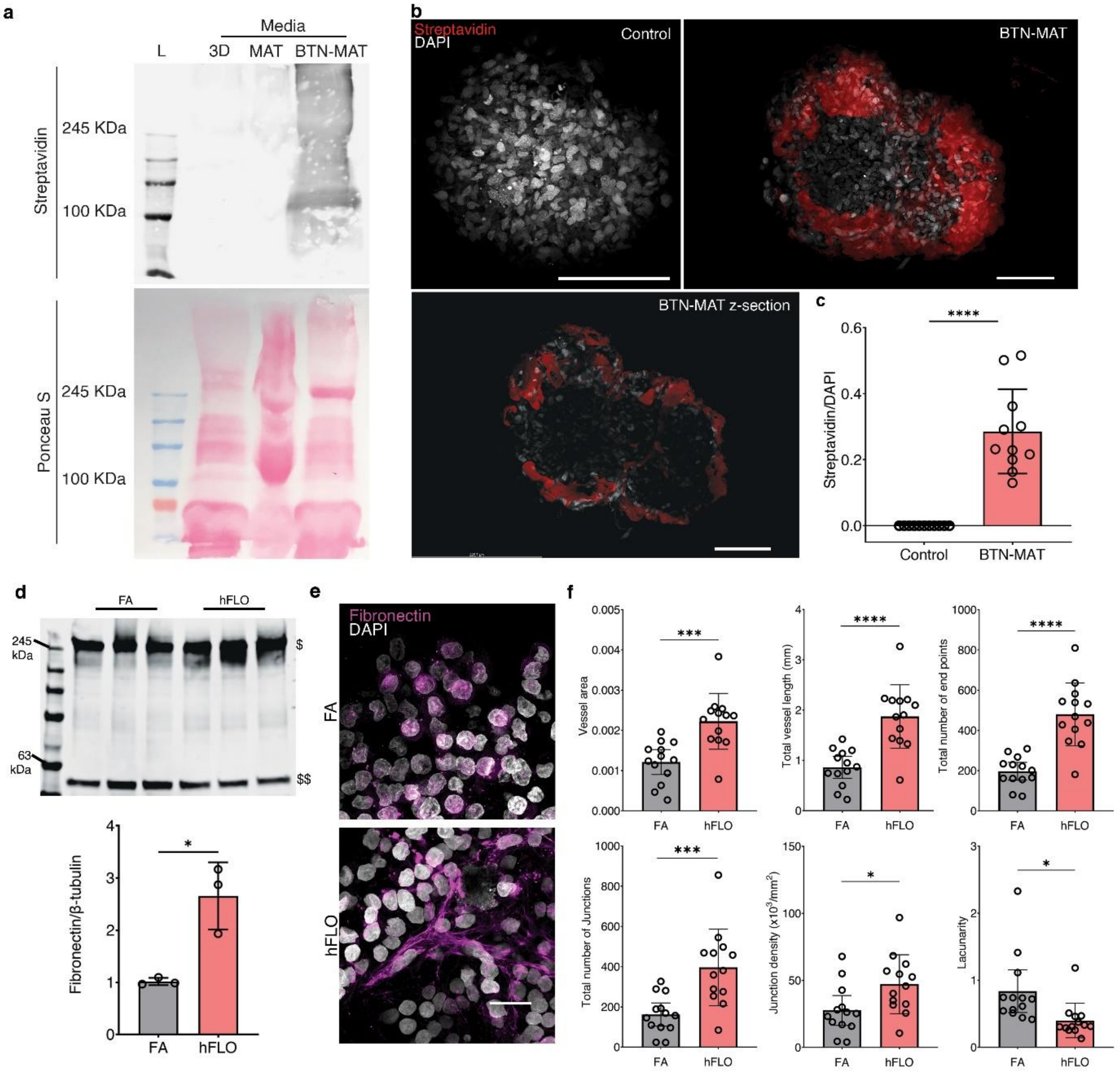


**Supplementary Fig. 6: Matrigel deposition in the aggregates.**

**a**, Immunoblot and ponceau staining using 3D media, hFLO media, and biotinylated Matrigel media (BTN-MAT). **b**, Representative fluorescence confocal images (maximum projection, Top; single z-section, Bottom) of hFLO at day 3 with normal matrigel media (Control) and biotinylated matrigel media (BTN-MAT). Scale bars, 100 µm. **c,** Volumetric quantification of biotinylated Matrigel localization in the aggregate normalized to nuclear volume (DAPI). 11 independently grown aggregates were used in the study. ****$P$<0.0001 determined using two-tailed Welch's t-test. **d,** Immunoblot using whole aggregate lysates probing for fibronectin ($). β-tubulin ($$) was used as loading control. Semi-quantitative analysis of the blot were normalized to β-tubulin. *$P$<0.05 determined using two-tailed Welch's t-test. **e,** Representative fluorescence confocal images (maximum projection) of FA and hFLO triculture aggregates at day 14 of culture immunostained for fibronectin. Scale bars, 20 µm. **f**, Quantification of fibronectin networks using AngioTool. 13 fields of view from 3 independently grown aggregates were used. Means, individual values, and standard deviations are shown. *$P$<0.05; ***$P$<0.001; ****$P$<0.0001 determined using two-tailed Welch's t-test.

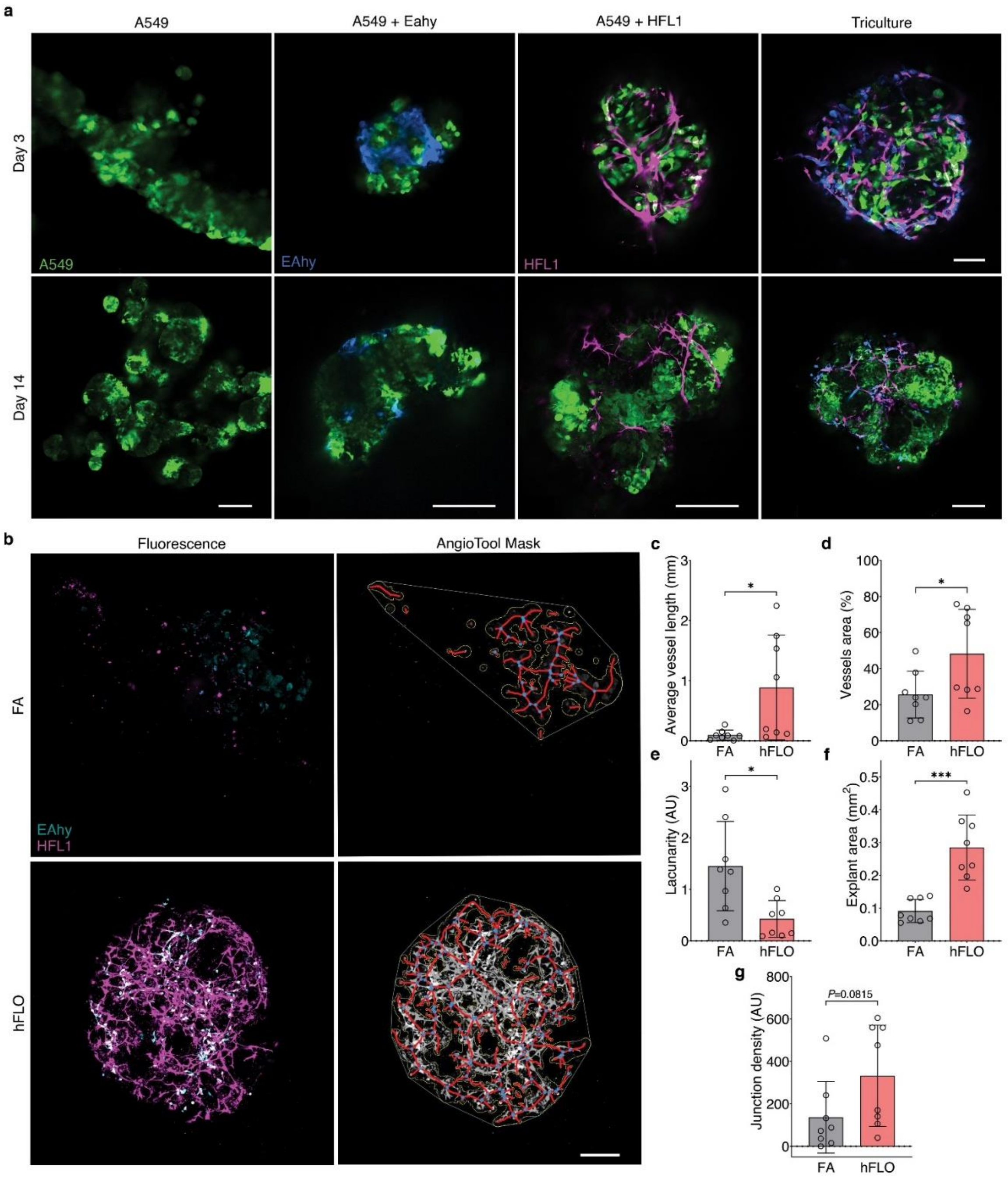


**Supplementary Fig. 7: FA versus hFLO AngioTool.**

**a,** Fluorescence confocal microscopy images showing the effects of co-culture to EAhy network formation in suspension. Co-culture with HFL1 is required for EAhy network formation. Scale bars, 100 µm for Day 3; 200 µm for Day 14. **b**, Representative maximum projection images acquired using confocal microscopy of FA and hFLO aggregates, filtered to show only mCherry (HFL1) and Azurite (EAhy) fluorescence. Vascular networks identified by AngioTool shown to the right. Scale bar, 100 µm. **c-g,** AngioTool quantification of vascular networks. In **d,** higher levels indicate more heterogeneity in the distribution of vascular networks. n = 8 aggregates for each treatment. Means, individual values, and standard deviations are shown. ns, not significant; **P*<0.05; ****P*<0.001; Welch's t-test.

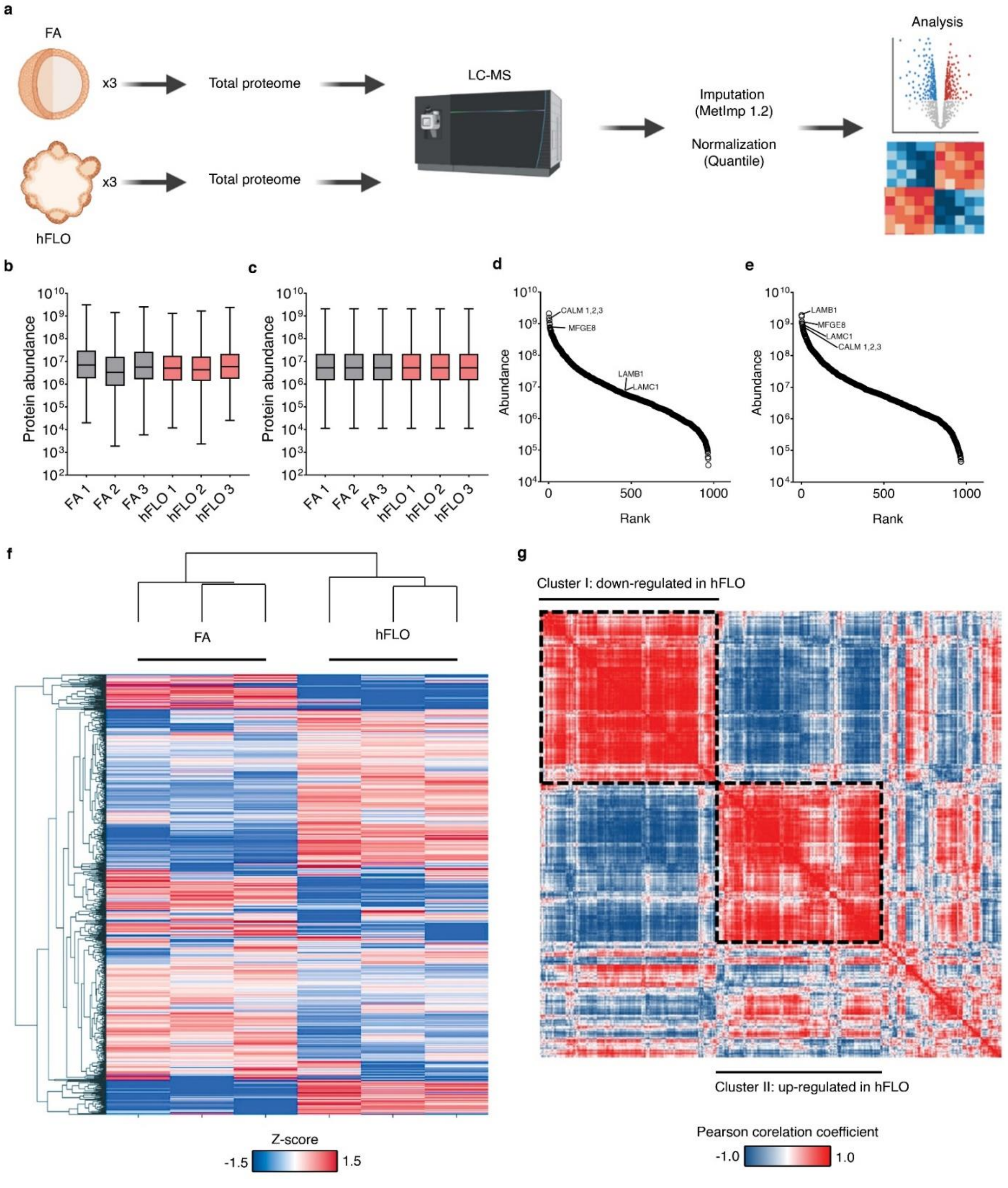


**Supplementary Fig. 8: Proteomic data processing**

**a**, Workflow for the proteomic profiling of FA and hFLO aggregates. Three independent samples were used for FA and hFLO; total proteome was extracted and run using a liquid chromatography-mass spectrometry (LC-MS). Proteomic data were input with MetImp and were quantile normalized. **b**, Boxplot of protein abundance (area) from all protein identified per sample before quantile normalization. **c**, After quantile normalization. **d**, Proteins in FA ranked based on abundance. **e**, Proteins in hFLO ranked based on abundance. Some highly abundant proteins are basement membrane proteins such as LAMB1 and LAMC1. **f**, Cluster analysis across samples and proteins was performed using Euclidean distance based on the protein z-scores. **g**, Correlation map of all detected proteins indicating Euclidean distance between proteins. Pearson's correlation coefficient was used and hierarchical clustering was based on Euclidean distance. Black dashed line indicates major clusters found.

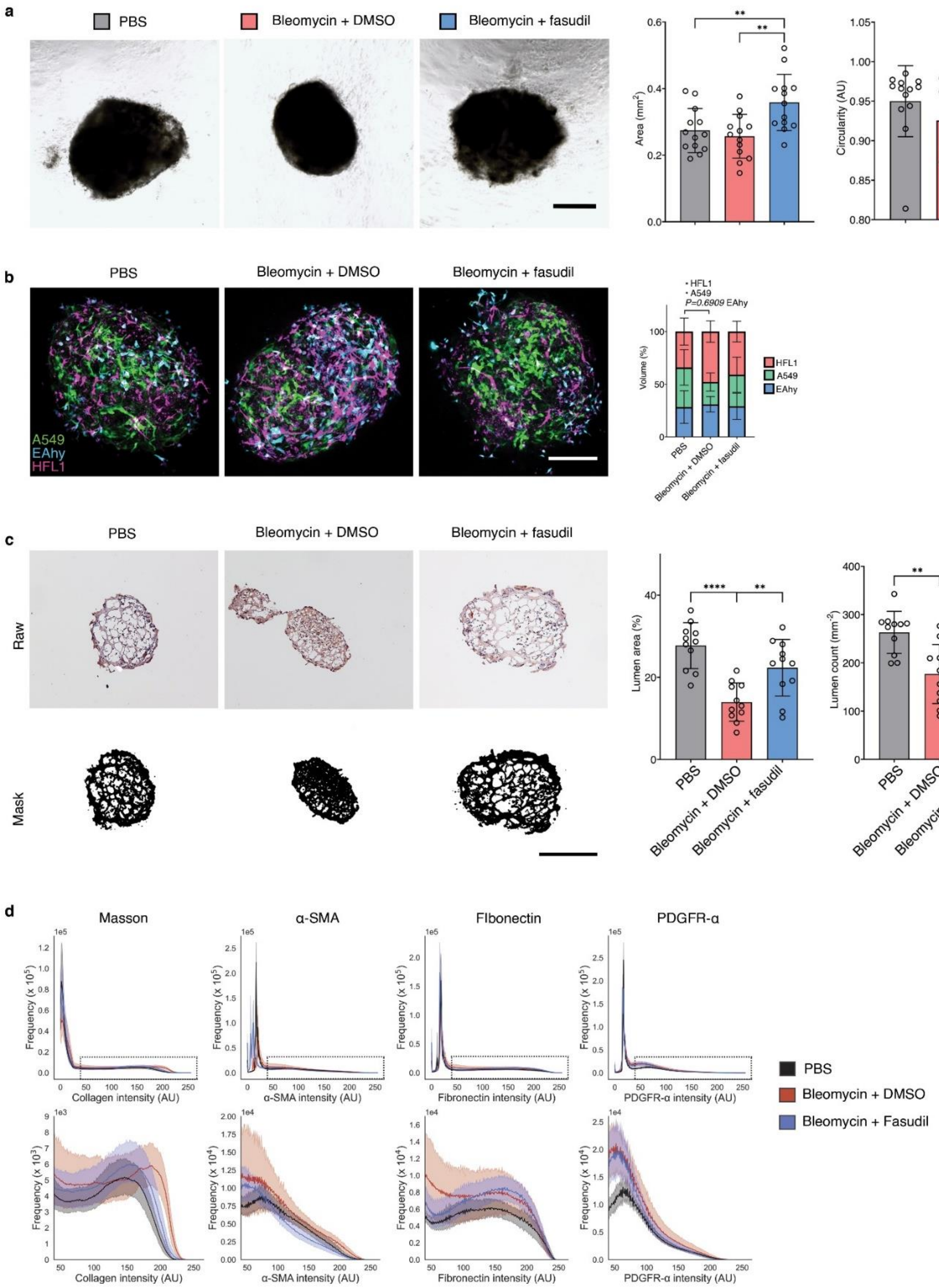


**Supplementary Fig. 9: Lumen analysis in fibrotic hFLO.**

**a**, Representative bright-field images of day 14 hFLO fibrotic model. Scale bar, 300 µm. Bar graphs for aggregate size and circularity. Means, standard deviation, and individual points are shown. We used n=13 independent cell aggregates. ***P*<0.01 determined using One-way ANOVA with Welch's correction. **b**, Live 3D fluorescence confocal imaging of the aggregates (maximum projection). Scale bar, 200 µm. Cell line volumetric analysis among treatment groups. *P<0.05 determined using One-way ANOVA with Welch's correction. **c**, H-DAB stained aggregates, showing the raw images (Top) and the images after ImageJ processing (Bottom). Scale bar, 250 µm. Quantification of lumen characteristics using ImageJ show means, individual values, and standard deviations are shown. n=11 independent aggregates; **P*<0.05, ***P*<0.01, *****P*<0.0001 determined using One-way ANOVA with Welch's correction. **d**, Whole spectrum histogram of intensity of pro-fibrotic markers collagen deposition (Masson), α-SMA, fibronectin, and PDGFR-α (Top). Zoom in on intensities ≥50AU (Bottom). We used n=9, n=11, n=9, and n=9 independent aggregates for each condition, respectively. Means and 95% confidence intervals are shown.

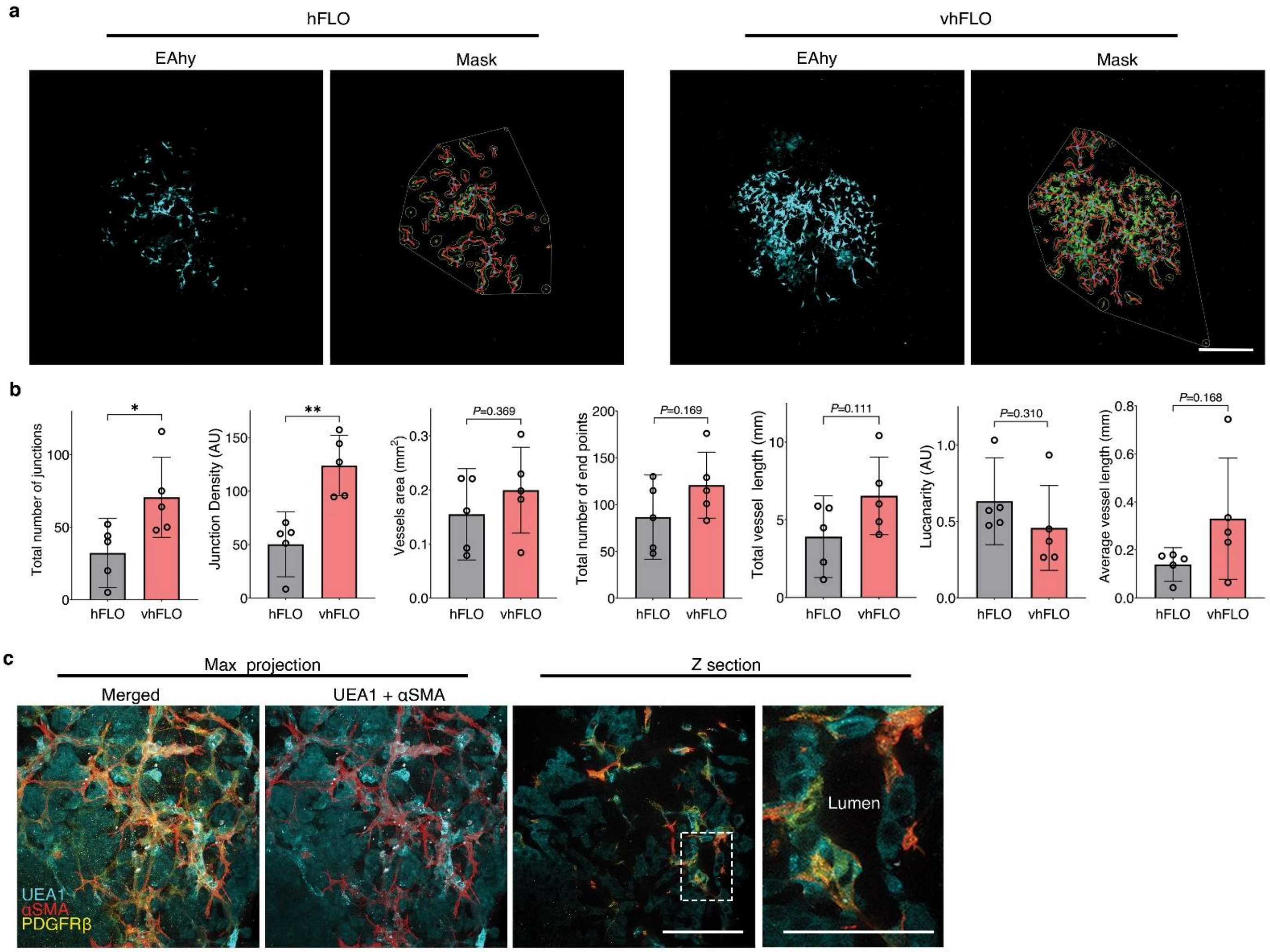


**Supplementary Fig. 10: Vascular cells in vhFLO.**

**a**, Live fluorescence confocal images (maximum projection) of EAhy cells in day 14 hFLO and vhFLO and AngioTool masks. Scale bar, 300 µm. **b**, Quantification of endothelial networks using AngioTool. We used 5 independent cell aggregates. Graphs show means, individual values, and standard deviation. **P*<0.05, ***P*<0.01 determined using Two-tailed Welch's t-test. **c**, Fluorescence confocal microscopy *maximum projection) using day 14 hFLO showing vascular and perivascular cells stained using UEA1, αSMA, and PDGFRβ. Single z-section image shows luminal morphology in hFLO vasculature. Scale bar, 100 µm; zoomed image scale bar, 50 µm.

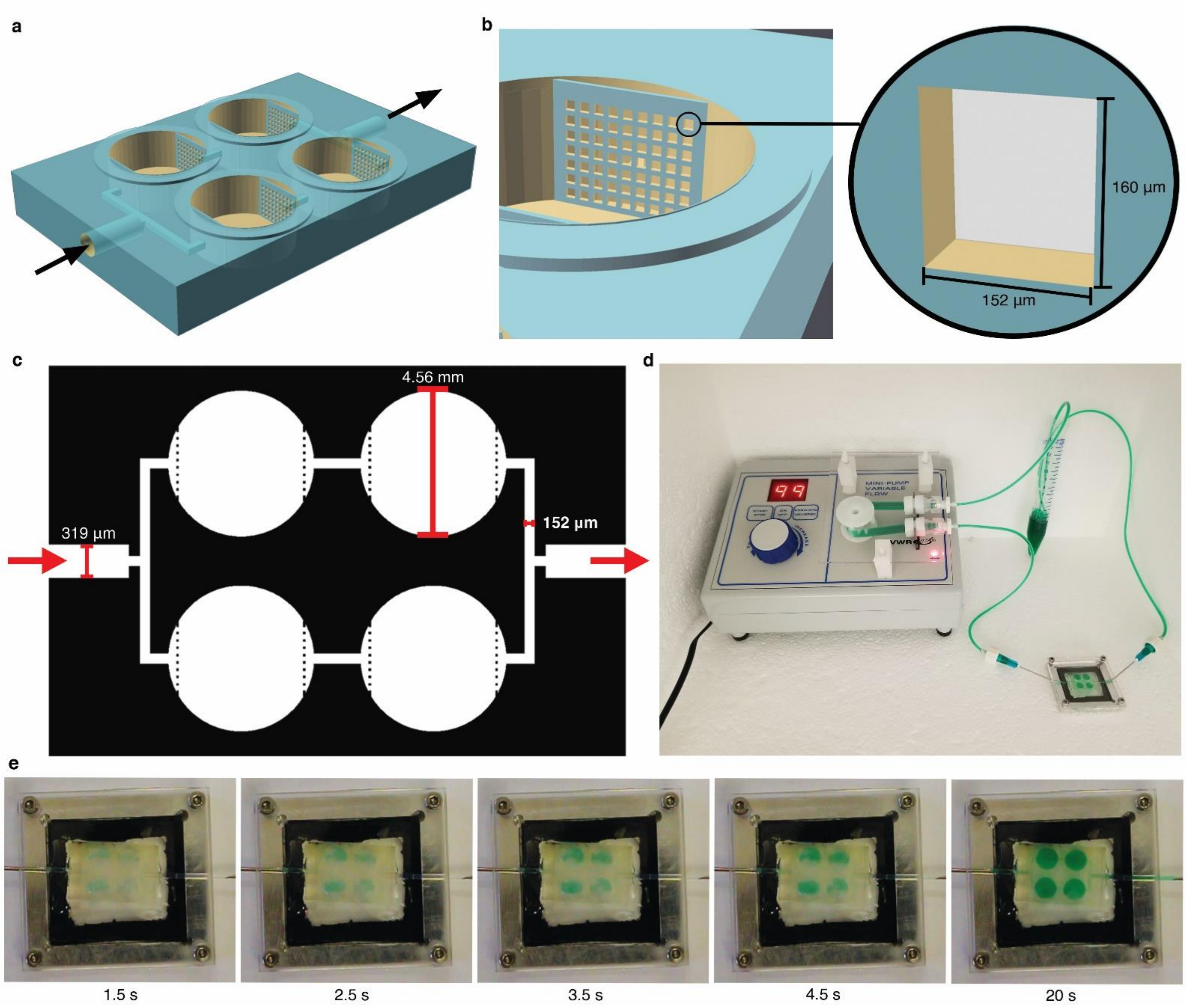


**Supplementary Fig. 11: Perfusion setup.**

**a**, CAD render of the 3D printed PEGDA resin perfusion chip; arrows indicate fluidic flow. **b**, Zoomed image showing a micro-grate and its dimensions. **c**, 2D layout of the chip and dimensions of notable features; micro-grates are represented by dotted lines. **d**, Setup of the perfusion chip-pump system. **e**, Time lapse showing even perfusion of dye through both channels over time; perfusion rate is set at 1 mL $min^{-1}$ $well^{-1}$.

### SUPPLEMENTARY DATA

**Table S1 Cell composition of lung and hFLO triculture**

| Cell type | Whole lung | Triculture | Per 1.0 x $10^4$ cell aggregate |
|---|---|---|---|
| Epithelial cells* | 24% of cells | 20% of cells | 2.0 x $10^3$ |
| Endothelial cells** | 29% of cells | 40% of cells | 4.0 x $10^3$ |
| Interstitial cells*** | 36% of cells | 40% of cells | 4.0 x $10^3$ |
| Other cells | 10% of cells | - | - |

*This includes both Type I and Type II alveolar cells for whole lung. In hFLO, only A549 cells are used.
**In hFLO, endothelial cells are modeled with the EAhy cell line.
***In hFLO, interstitial cells are modeled with the HFL1 cell line.

**Table S4 Media formulations and notes**

| Media | Contents | Notes |
|---|---|---|
| 2D growth media | DMEM/F12<br>+10% Fetal bovine serum (FBS)<br>+Anti-biotic/ anti-mycotic (anti-anti) (Carson)<br>+ Plasmocin prophylactic (Invivogen) | For 2D culture of A549, HFL1, and LentiX |
| 3D growth media | DMEM/F12<br>+10% Fetal bovine serum (FBS)<br>+Anti-biotic/ anti-mycotic (anti-anti) (Carson)<br>+ Plasmocin<br>+ 0.024% methylcellulose (Sigma) | For 3D culture of monoculture and triculture cell aggregates. Metylcellulose enhances aggregate compaction. |
| hFLO media | DMEM/F12<br>+ 10% Fetal bovine serum (FBS)<br>+ Anti-biotic/ anti-mycotic (anti-anti) (Carson)<br>+ Plasmocin prophylactic (Invivogen)<br>+ 0.024% methylcellulose (Sigma)<br>+ 300 µg $mL^{-1}$ Matrigel GFR | The minimum gelling concentration of Matrigel is 3 mg $mL^{-1}$ according to the manufacturer*. Matrigel concentration in the hFLO media keeps the Matrigel proteins solubilized in media. |
| Vascularization media (vhFLO media) | DMEM/F12<br>+ 10% Fetal bovine serum (FBS)<br>+ Anti-biotic/ anti-mycotic (anti-anti) (Carson)<br>+ Plasmocin prophylactic (Invivogen)<br>+ 0.024% methylcellulose (Sigma)<br>+ 300 µg $mL^{-1}$ Matrigel GFR.<br>+ 60 ng $mL^{-1}$ FGF2<br>+ 60 ng $mL^{-1}$ VEGF-165 | FGF2 and VEGF are pro-vascular factors. |

*See Corning® Matrigel® Matrix Frequently Asked Questions (CLS-DL-CC-026)

**Table S5 Dimensions of 3D printed millifluidic perfusion device**

| Part | Dimension | Value |
|---|---|---|
| Bulk | Length | 18.00 mm |
| | Width | 12.16 mm |
| | Height | 2.50 mm |
| Input/ Output channels | Width | 319 µm |
| | Height | 300 µm |
| Wells | Diameter | 4.56 mm |
| | Depth | 0.50 mm |

**Table S6 Primary antibodies and Stains**

| Antibody (Anti-) | Figures* | Source | Catalog number | Dilution (Application**) |
|---|---|---|---|---|
| Fibronectin | F5, EF5, EF8 | ProteinTech | 66042-1-Ig | 1:200 (IF)<br>1:200 (IHC-HRP) |
| Vimentin | F1 | Cell Signaling Technology | D21H3 | 1:100 (IF) |
| PDGFRβ | F3, F6, EF9 | Cell Signaling Technology | 3169 | 1:100 (IF)<br>1:100(FC) |
| PDGFR α | F5, EF8 | Cell Signaling Technology | 3174 | 1:1000 (IHC-HRP) |
| αSMA | F3, F5, F6, EF8, EF9 | Cell Signaling Technology | 48938 | 1:200 (IF)<br>1:200 (FC)<br>1:200 (IHC-HRP) |
| ECadherin | EF1 | Cell Signaling Technology | 3195 | 1:200 (IF) |
| NCadherin | EF1 | Thermo Fisher Scientific | 33-3900 | 1:150 (IF)<br>1:150 (FACS) |
| EPCAM | F1 | Developmental Studies Hybridoma Bank | G8.8 | 1:50 (IF) |
| Pro-SPC | F1 | Abcam | ab40879 | 1:250 (IF) |
| PDPN | F1 | Thermo Fisher Scientific | 14-5381-82 | 1:100 (IF) |
| Desmin | F3 | Invitrogen | 14-9647-82 | 1:100 (IF)<br>1:100 (FC) |
| Rhodamine UEA1 | F3, F6, EF9 | Vector Laboratories | RL-1062-2 | 1:100 (IF) |
| Cleaved Caspase-3 (Asp175) | F2, | Cell Signaling Technology | 9661 | 1:800 (MIFC) |
| Streptavadin -CF™633 | EF5 | Biotium | 29037 | 1:500 (IF) |

*F=Figure; EF=Extended Figure

**IF=Immunofluorescence; FC=Flow Cytometry; FACS=Flow-Activated Cell Sorting; MIFC=Multispectral Imaging Flow Cytometry; IHC-HRP=Immunohistochemistry-Horseradish Peroxidase
